# Two Parallel Pathways Mediate Olfactory-Driven Backward Locomotion

**DOI:** 10.1101/2020.11.23.393819

**Authors:** Shai Israel, Eyal Rozenfeld, Denise Weber, Wolf Huetteroth, Moshe Parnas

## Abstract

Although animals switch to backward walking upon sensing an obstacle or danger in their path, the initiation and execution of backward locomotion is poorly understood. The discovery of Moonwalker Descending Neurons (MDNs), made *Drosophila* useful to study neural circuits underlying backward locomotion. MDNs were demonstrated to receive visual and mechanosensory inputs. However, whether other modalities converge onto MDNs and what are the neural circuits activating MDNs are unknown. We show that aversive but not appetitive olfactory input triggers MDN-mediated backward locomotion. We identify in each hemisphere, a single Moonwalker Subesophageal Zone neuron (MooSEZ), which triggers backward locomotion. MooSEZs act both upstream and in parallel to MDNs. Surprisingly, MooSEZs also respond mostly to aversive odor. Contrary to MDNs, blocking MooSEZs activity has little effect on odor-evoked backward locomotion. Thus, this work reveals another important modality input to MDNs in addition to a novel olfactory pathway and MDN-independent backward locomotion pathway.

## Introduction

Walking is a fundamental behavioral feature of many terrestrial organisms, allowing them to respond adaptably to internal and environmental stimuli. In order to support motor-based behaviors, and specifically rhythmic activities such as walking, the nervous system needs to perform a sequential multi-step process: collect relevant sensory information, integrate incoming information with internal-state status, and efficiently execute the selected motor program. Although the nervous system of invertebrates is compact in comparison to that of vertebrates, their walking repertoire is highly diverse and complex^1^. Thus, insects’ nervous systems can serve as an experimental model to study the neuronal mechanisms underlying coordinated motor activity. In particular, the fruit fly seems to be an ideal choice, as individual cells within its brain can be precisely targeted and easily manipulated using genetic techniques^2^.

Several decades of research have led to substantial knowledge about basic motor pattern generation in insects, such as reflex-activated leg movements^3,4^, jumping^5^, wing movement in flying^6,7^, and leg stepping in walking^8,9^. Recently, more sophisticated behaviors were examined, such as coordination of antennae and leg movement during obstacle negotiation^10,11^, gap crossing^12^, grooming^13^, chemotaxis orientation^14^ and courtship behavior^15,16^. Although most land animals switch to backward walking upon sensing an obstacle or danger in their path, the initiation and execution of backward walking is still poorly understood^1^.

By exploiting the powerful genetic toolkit available for fruit flies^17^, a specific cluster of neurons, Moonwalker Descending Neurons (MDNs), which triggers backward walking was identified^18^. In addition, it was shown that a population of neurons in the fly’s visual system induce backward walking via MDNs through indirect synaptic connections^19^ and that ascending mechanosensory neurons in the ventral nerve cord (VNC) activate MDNs to mediate touch-evoked backward walking^20^. However, other sensory pathways such as gustation, thermoreception and in particular olfaction, a major sensory input on which insects rely heavily^21^, have not been associated with MDN mediated backward walking. Furthermore, characterization of the neuronal circuits which functionally activate or modulate MDNs and thus backward walking is currently lacking.

In the case of flies, odors are sensed by first-order olfactory sensory neurons (OSNs) expressing either odorant receptors (ORs) or ionotropic receptors (IRs)^22–26^. There are 51 types of OSNs defined by the type of OR or IR they express^27^. OSNs project to the antennal lobe (AL), where the axons of each type of OSN target a single glomerulus^28–30^. Second order projection neurons (PNs) send their dendrites to a single glomerulus and project to two higher brain regions: the mushroom body (MB), where associative olfactory memories occur and the lateral horn (LH)^31^.

In the present study we show that aversive but not appetitive olfactory input can trigger backward locomotion mediated by MDNs. In addition, we identify a single neuron in each hemisphere originating in the subesophageal zone (SEZ) which triggers backward walking. We name this neuron Moonwalker SEZ (MooSEZ). MooSEZs are functionally connected to MDNs but can also trigger backward walking in an MDN-independent manner. Surprisingly, we find that MooSEZs respond to aversive but not to appetitive odors. Whereas blocking MDNs activity eliminates odor-driven backward walking, blocking MooSEZ activity has little effect. This suggests that MDNs are both sufficient and necessary whereas MooSEZ are sufficient but not necessary for olfactory-driven backward locomotion. Thus, this work reveals another important modality input to MDNs in addition to a novel olfactory pathway and MDN independent backward locomotion pathway.

## Results

### Olfactory input triggers MDN-dependent backward walking

Recently it was demonstrated that visual input to MDNs generates backward walking^19^. It was suggested that this backward locomotion is part of a repertoire of escape responses to a visual threat. Similar to visual inputs, olfactory cues can also signal threat^32,33^. We first examined whether 2-butanone, a strongly aversive odor^34^, can trigger backward walking. To this end we placed flies in an open field arena and examined their responses when exposed to 2-butanone. While in most flies exposure to 2-butanone elicited turning and forward walking, in ~25% of the flies turning was combined with a pronounced backward walking component (Figure 1A-C). To verify that the observed backward walking response is indeed an odor-specific effect we used the attractant odor apple cider vinegar^34^ (ACV). Following exposure to ACV we observed negligible retreat response across tested flies (Figure 1A, C). Yet, the backward walking responses evoked by 2-butanone in the open field arena were rather transient, consisting of only a number of retreat steps coupled to odor presentation. We therefore sought to enhance the backward walking phenotype and repeated the experiment in a linear chamber assay system that was shown to effectively restrict flies’ lateral and rotational movements (Figure 1D, left). Using this apparatus, both 2-butanone and ACV efficiently induced backward walking responses among flies. However, strong and robust backward walking was observed only for 2-butanone but not for ACV (Figure 1D right, 1E). Since ACV valence is known to depend on the animals’ satiation state but 2-butanone remains highly aversive even in starved flies^34^, we repeated the linear chamber experiment using starved flies. Indeed, backward walking following exposure to ACV was now abolished, whereas 2-butanone still elicited strong backward walking (Figure 1F and Movie S1).

**Figure 1:**
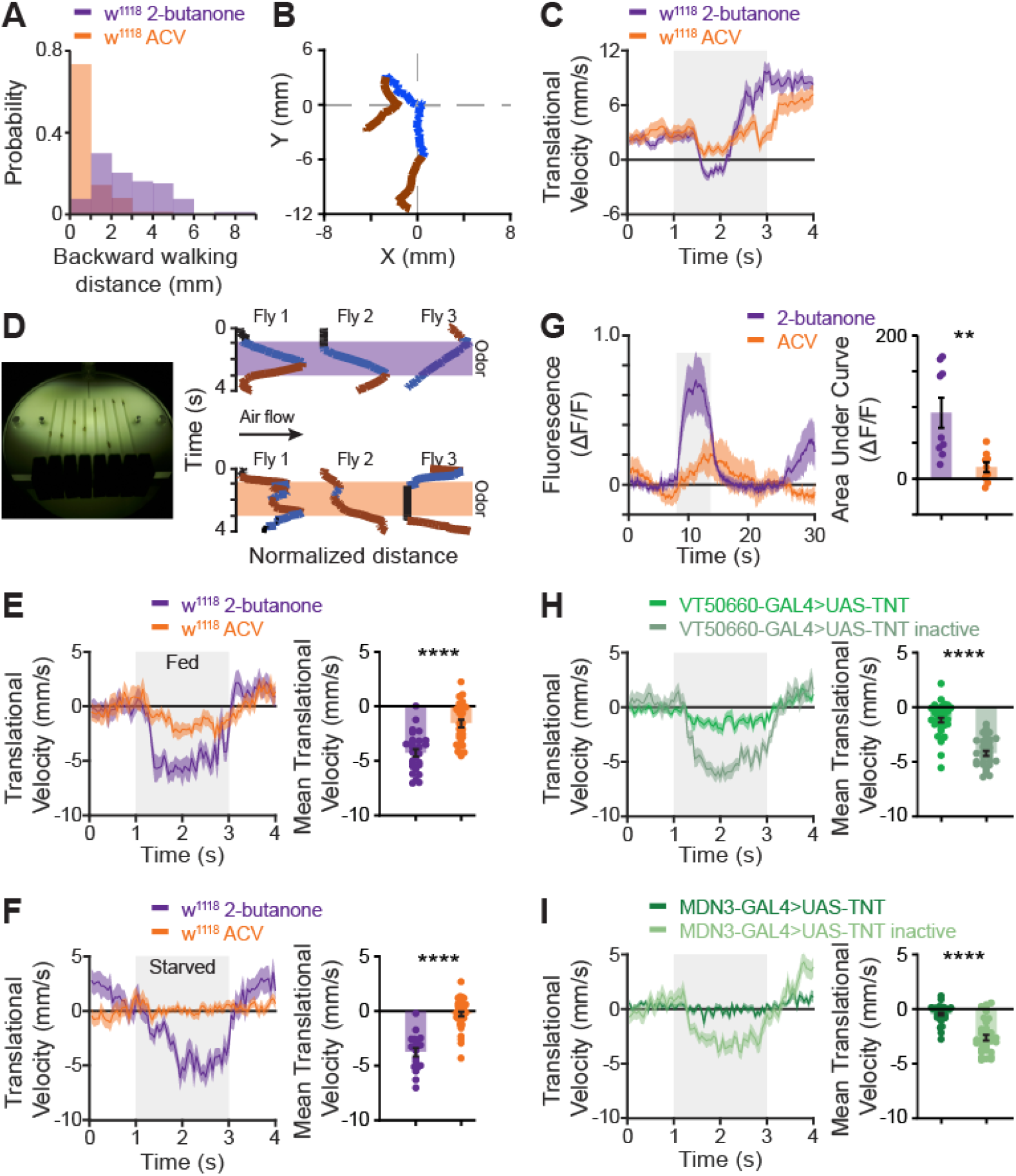
Odor-induced backward walking is mediated by MDNs. **(A)** The probability of observing backward locomotion in response to an odor pulse in an open arena. For the appetitive odor ACV no backward walking was observed whereas backward walking was observed for the aversive odor 2-butanone approximately. **(B)** Examples of two fly trajectories following application of 2-butanone. Flies responded in a transient backward locomotion. Blue and brown lines designate backward and forward locomotion respectively. **(C)** Translational velocity ± SEM (shading) following application of 2-butanone (dark purple) and ACV (orange). The odor pulse is labelled in grey. **(D)** *Left,* the linear chamber behavior apparatus. *Right,* Examples of flies’ trajectory in the linear chambers. Black denotes stationary flies. Blue denotes backward walking, brown forward motion, and black stalling. 2-butanone is labelled in purple, ACV is labelled in orange. **(E, F)** *Left*, translational velocity ± SEM (shading) following application of 2-butanone (dark purple), and ACV (orange) in fed (E) or starved (F) flies. The odor is labelled in grey. A clear backward motion is observed following 2-butanone but not following ACV. *Right*, mean translational velocity during the entire odor pulse obtained from traces on the left. A significant difference is observed between the backward walking velocity generated by 2-butanone and that elicited by ACV (fed: 30≥n≥28, starved: 41≥n≥19 **** p < 0.0001, Mann-Whitney test). **(G)** *Left,* averaged traces ± SEM (shading) of odor responses (as indicated, odor pulse is labelled in grey). VT50660-GAL4 was used to drive GcaMP6f in MDNs. *Right,* area under the curve of ΔF/F during odor response for the traces presented in the *left* panel. The aversive odor 2-butanone has a stronger odor response compared to the appetitive odor ACV. (n = 9 flies, ** p < 0.01, Two-sample t-test). **(H, I)** *Left,* translational velocity ± SEM (shading) following application of 2-butanone for flies expressing TNT under the control of the broad MDN driver line VT50660-GAL4 (H, green) or under the control of the narrow split GAL4 driver line MDN3-GAL4 (I, dark green). Odor pulse is labelled in grey. The inactive TNT is labeled by light green. In both cases the expression of TNT dramatically decreases backward walking. *Right,* mean translational velocity during the entire odor pulse obtained from traces on the left. A significant difference is observed between the backward walking velocity generated by 2-butanone when TNT is expressed compared to when the inactive TNT is expressed. (MDN3: 37≥n≥35, VT50660: 40≥n≥32, **** p < 0.0001, Mann-Whitney test).

As MDNs were demonstrated to be causally link to backward walking^18^, we examined whether an odor pulse can trigger neuronal responses in MDNs. We expressed GCaMP6f^35^ in MDNs using the broad MDN driver line VT50660-GAL4 and performed 2-photon *in vivo* Ca^2+^ imaging, focusing on the dendritic arbors of the MDNs. As expected from the results above, MDNs showed strong responses to application of 2-butanone but not to application of ACV (Figure 1G). To examine whether MDNs are necessary for olfactory-driven backward locomotion, we expressed in MDNs the tetanus toxin light chain (TNT), which is an inhibitor of synaptic transmission^36^, using either the broad MDN driver, VT050660-GAL4, or the more specific split-GAL4, MDN3-GAL4^18^. Expression of TNT in MDNs abolished odor-driven backward locomotion. Expression of the inactive tetanus toxin (TNT-inactive) had no effect on the odor-elicited backward walking (Figure 1H, I). Taken together, we found a backward walking response specific to aversive odor that requires MDNs.

### Optogenetic activation of GH146^II^, NP225 and NP5288-GAL4 triggers backward walking

The results above suggest that an olfactory input can trigger backward walking via MDNs. In an attempt to identify the underlying neurons responsible for this effect, we screened 43 driver lines covering different populations of olfactory neurons by acute optogenetic activation with ChR2-XXM^37^ or CsChrimson^38^. We first verified that we can optogenetically trigger and identify backward walking by activating MDNs. Activation of MDNs using both the broad and narrow MDN driver lines indeed elicited sustained backward walking, as previously observed^18^ (Figure S1). Out of the 43 drivers examined, we observed robust and prolonged backward walking in three broad driver lines which cover most of PNs: GH146^II^-GAL4 (GH146-GAL4 on the second chromosome), NP5288-GAL4 and NP225-GAL4 (Figure 2A, B and Movie S2). Importantly, MDNs are not labelled by these lines (Figure S2A and S3). Surprisingly, optogenetic activation of GH146-GAL4 on the X chromosome (GH146^X^) did not elicit backward locomotion (Figure 2A, B). In some driver lines, optogenetic activation resulted in a seizure-like behavior and thus these driver lines could not be examined (Figure 2A). We also characterized the sensitivity of the elicited backward walking to the light stimulus intensity. As expected, activation using blue light (470 nm) was intensity-sensitive and ranged from eliciting forward motion at low intensities to eliciting backward walking at high intensities (Figure S2B). Surprisingly, we found strong and efficient activation of backward walking at all tested red light (617 nm) intensities (Figure S2B).

**Figure 2:**
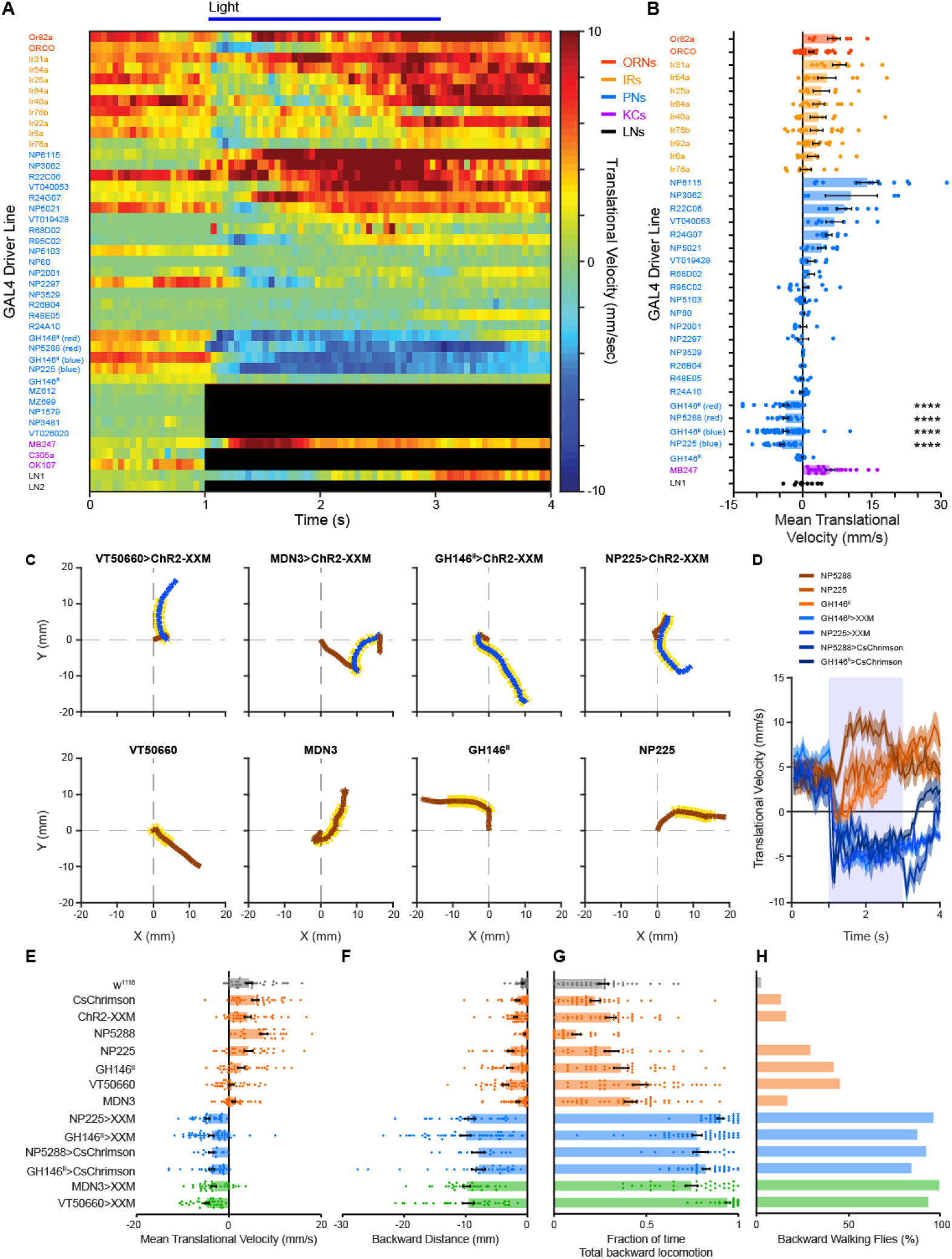
GH146^II^-, NP225- and NP5288-GAL4 induce similar backward walking as MDNs. **(A)** Translational velocity matrix of different GAL4 driver lines (as designated) expressing ChR2-XXM (or CsChrimson where designated, red). A light pulse (horizontal blue line) was given between 1 and 3 seconds. In some cases light activation of ChR2-XXM resulted in a seizure like behavior which did not allow assessing walking velocity. Such cases are labeled in black. **(B)** Mean translational velocity during the entire light pulse obtained from traces used to compose the matrix in A. A significant backward locomotion is observed for GH146^II^-, NP225- and NP5288-GAL4 irrespective of the channel used for activation. (50≥n≥4, **** P < 0.0001, One sample t-test against zero (one-tailed) for each driver line, followed by False Discovery Rate correction). **(C)** Examples of single fly walking trajectories for MDNs, GH146^II^-, NP225-GAL4 driving ChR2-XXM and parental controls. Light pulse is designated by yellow, blue denotes backward walking and brown forward motion. Similar backward walking pattern can be observed for all flies which is lacking in parental controls. **(D)** Translational velocity ± SEM (shading) for GH146^II^-, NP225-, or NP5288-GAL4 driving either UAS-ChR2-XXM or UAS-CsChrimson as designated. Sustained backward walking is observed throughout the two second light pulse (labelled by light blue). **(E-H)** Analysis of the following parameters during the 2s light pulse: (E) mean translational velocity, (F) backward distance covered by the flies, (G) fraction of time in which flies’ translational velocity was negative, and (H) the percentage of flies that covered a minimal 3 mm backward walking distance for MDN driver lines (green), GH146^II^-, NP225-, or NP5288-GAL4 driver lines (blue), parental controls (brown), and *wt* flies (grey). For all computed parameters, GH146^II^-, NP225-, and NP5288-GAL4 driver lines resemble MDN driver lines. (50≥n≥14, p < 0.0001 for all comparisons to respective parental controls and *wt* except for mean translational velocity, MDN3 p < 0.05, mean backward distance, GH146^II^>CsChrimson p < 0.01, fraction of time of total backward locomotion, MDN3 p < 0.01. Kruskal – Wallis test followed by Dunn’s post-hoc test. Percentage of backward walking flies: p < 0.001 for all comparisons to respective parental controls and wt, chi-squared test, followed by false discovery rate correction).

Activation of MDNs elicits robust and sustained backward walking with a weak angular change (Figure 2C and S1). In contrast, activation of the visual and mechanosensory pathways that activate MDNs elicit pronounced but transient backward walking with a strong turning component^19,20^. We replicated these response dynamics by optogenetically activating LC16-1-GAL4, which drives expression in a distinct class of optic interneurons connecting the lobula to an optic glomerulus in the protocerebrum, and TLA-GAL4, which drives expression in TwoLumps Ascending Neurons (Figure S2C, D). Interestingly, the backward locomotion following 2 second optogenetic activation of GH146^II^-, NP225, and NP5288-GAL4 was distinctively similar to the persistent, straight motor response evoked by optogenetic activation of MDNs, as opposed to the transient and curved responses generated for LC16-1-GAL4 and TLA-GAL4 optogenetic activation (Figure 2A, C, D). We also compared a number of parameters describing the efficiency of activation of backward walking and found no significant differences between MDN activation using MDN3 and VT50660 and the three GAL4 driver lines, GH146^II^, NP225, and NP5288 (Figures 2E-H).

### Backward locomotion induced by GH146^II^ has MDN-dependent and -independent components

We have found that neurons labelled by GH146^II^ drive backward locomotion, whereas those labelled by GH146^X^ do not. This suggests that the GH146-induced backward locomotion is not mediated by PNs covered by both GH146 driver lines. We therefore examined whether the GH146-evoked backward locomotion is mediated by MDNs. To this end, we asked whether GH146^II^-GAL4 neurons could trigger neuronal responses in MDNs. We optogenetically activated *ex vivo* the GH146^II^-GAL4 neurons with CsChrimson and examined calcium transients in the dendritic arbors of MDNs using the genetically encoded Ca^2+^ indicator GCaMP6m^35^. Indeed we observed robust responses in MDNs after optogenetic activation of GH146^II^-GAL4 neurons, although only at high light intensity. Surprisingly, illuminating flies with low light intensity did not elicit calcium transients in MDNs’ dendrites, though flies did exhibit backward walking in response to this light stimulation (Figure 3A, S2B). The differential effect of light intensity on MDNs calcium responses may suggest that GH146^II^-GAL4-induced backward walking phenotype is not exclusively dependent on MDNs functionality.

**Figure 3:**
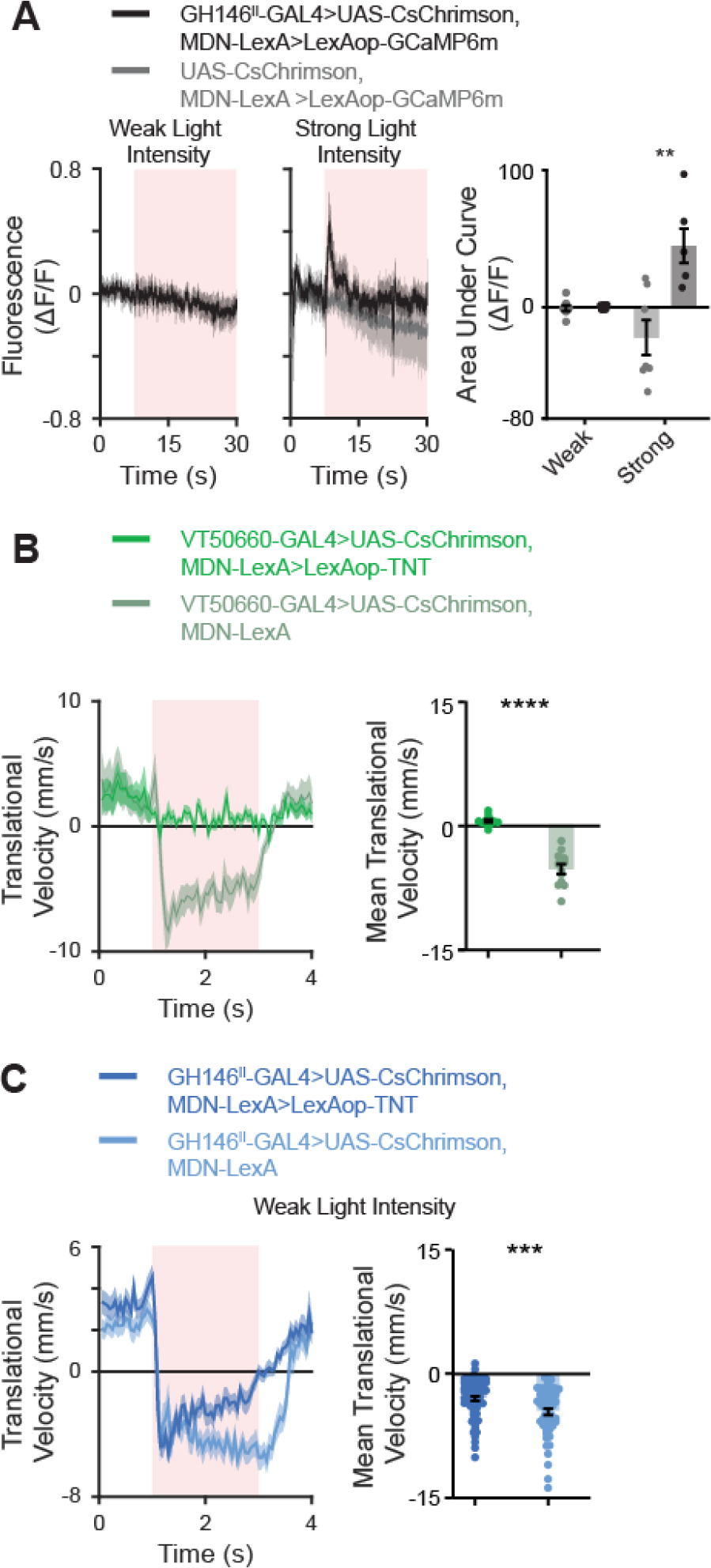
GH146^II^ neurons mediating backward locomotion are both MDN-dependent and – independent. **(A)** *Left,* averaged traces ± SEM (shading) of responses measured in MDNs following optogenetic activation of GH16 neurons using weak or strong light intensities of 3.35 mW/cm^2^ and 19.43 mW/cm^2^ respectively as designated. For the experimental group, labelled in black, the MDN-LexA driver line was used to drive LexAop-GCaMP6m and GH146^II^-GAL4 to drive UAS-CsChrimson. As a control, labelled in grey, the same genotype was used except that the GH146^II^-GAL4 was omitted. Light pulse is labeled in light red. *Right,* Area under the curve of ΔF/F during the light pulse for the traces presented in the *left* panels. For weak light intensities and control flies no response was observed in MDNs. A significant response was observed at strong light intensities. (9≥n≥6, *** p < 0.01, Two-sample t-test). **(B)** *Left,* translational velocity ± SEM (shading) following optogenetic activation of MDNs using weak light pulse (3.35 mW/cm^2^) in the presence (dark green) or absence (light green) of TNT as designated. The VT50660-GAL4 driver was used to drive UAS-CsChrimson, and the VT44845-LexA (MDN-LexA) was used to drive LexAop-TNT (when required). Light pulse is labeled in light red. *Right*, mean translational velocity during the entire light pulse obtained from traces on the left. TNT in MDNs completely blocks the backward walking and a significant difference is observed between the backward walking velocities when TNT is expressed compared to when TNT is not expressed. (12≥n≥11, **** p < 0.0001, Mann-Whitney test). **(C)** *Left*, translational velocity ± SEM (shading) following optogenetic activation of GH146^II^ neurons using weak light pulse (3.35 mW/cm^2^) in the presence (dark blue) or absence (light blue) of TNT in MDNs as designated. The GH146^II^-GAL4 driver was used to drive UAS-CsChrimson and the VT44845-LexA driver was used to drive LexAop-TNT (when required). Light pulse is labeled in light red. *Right,* mean translational velocity during the entire light pulse obtained from traces on the left. Robust backward locomotion is generated by weak light pulses even though MDNs activity is blocked by TNT (64≥n≥58, *** p < 0.001, Mann-Whitney test).

We then examined whether MDNs are necessary for GH146^II^-GAL4-evoked backward locomotion. To test if backward walking induced by artificial GH146^II^-GAL4 activation requires the established MDN neural pathway or acts via a parallel pathway, we inactivated MDNs by expressing TNT using MDN-LexA and lexAoP-TNT^18^. As a positive control, the efficiency of the lexAop-TNT was confirmed by direct optogenetic activation of MDNs. TNT efficiently silenced MDNs and eliminated backward walking (Figure 3B and Movie S3). Silencing MDNs and activating GH146^II^-GAL4 neurons at high light intensity (28.27 mW/cm^2^) caused flies to lose balance and fall on their back, raising the possibility that MDNs are recruited at strong excitation of GH146^II^-GAL4 neurons to stabilize backward walking. Yet, at low light intensity (3.35 mW/cm^2^), blocking MDNs did not abolish backward locomotion elicited by optogenetic activation of GH146^II^ (Figure 3C and Movie S3). Thus, these results suggest that GH146^II^–GAL4 labels neurons which are either downstream to MDNs or participate in a parallel pathway to MDNs. Taken together, our results demonstrate that the GH146^II^-GAL4 neurons inducing backwards walking are upstream to MDNs, but that they also participate in an additional locomotion pathway either downstream or in parallel to MDNs.

### Anatomical analysis of GH146, NP225, and NP5288 GAL4 lines

To identify candidate neurons covered by GH146^II^-, NP225-, and NP5288-GAL4 driver lines (other than the olfactory pathway) we conducted a comprehensive anatomical analysis of these lines. All lines exhibit very similar expression pattern in both brain and thoracico-abdominal ganglia (TAG); Notable exceptions are the APL neuron in the mushroom body that is exclusively labelled by GH146^II^–GAL4 (Figure S3), as summarized previously^39^, and additional somata in the central brain and TAG labelled by NP5288-GAL4. The additional expression shared between all three lines includes two paired lateral somata and one pair of medioventral somata in the SEZ, as well as several paired neurons across TAG segments, including a paired neuron in the posterior TAG that ascends into the contralateral protocerebrum (Figure S3).

### GH146^II^-, NP225, and NP5288-GAL4 thoracico-abdominal ganglia neurons do not contribute to backward walking

The anatomical analysis revealed labeling in the thoracico-abdominal ganglia (TAG). Since we found MDN-independent backward locomotion pathway, it is possible that TAG neuron activity downstream of MDNs underlies the observed backward locomotion. We addressed this option using two approaches. In order to isolate TAG neurons from central brain neurons, flies were decapitated. Such chirurgical manipulation removes any central brain neurons, but maintains ascending and descending tracts, such as axons of MDNs. Decapitated flies were shown to maintain coordinated walking following the application of octopamine to the cervical connective^40^. We first verified that decapitated flies have the ability to perform backward walking. Indeed, optogenetically activating MDNs using ChR2-XXM still resulted in robust backward walking (Figure 4A, B and Movie S4). We then optogenetically activated GH146^II^-, NP225-, and NP5288-GAL4 covered neurons in decapitated flies. No backward walking was observed (Figure 4C, D). However, we did observe behavioral responses to the optogenetic activation of GH146^II^, NP225, NP5288 TAG neurons. Flies changed their posture by shifting their body weights to meso- and metathoracal legs, lifting their forelegs and moving them in an oscillatory fashion (Movie S4). Investigation of this behavior was out of the scope of the current study and therefore we did not further analyze it. Thus, optogenetic activation of GH146^II^, NP225, and NP5288 TAG neurons affects flies’ behavior but does not generate any backward walking. As expected, no behavioral response whatsoever was observed in decapitated *wt* flies or any of the parental controls (Figure 4B, D). The above results suggest that optogenetic activation of GH146^II^-, NP225-, and NP5288-GAL4 neurons triggers two behavioral responses: the first is the backward walking which probably depends on labeled neurons in the central brain, and the second is the shift in body posture and foreleg movements which involves labeled neurons in the TAG. We therefore revisited the experiments used for Figure 2 and indeed observed a combination of the two behavioral responses (Movie S2). We then used tsh-GAL80 to block expression of CsChrimson in TAG neurons^15,41^ driven by GH146^II^-, NP225-, and NP5288-GAL4, which abolished expression in most TAG neurons and the medioventral SEZ somata, but spared the lateral SEZ neurons (Figure 4E). As expected from the above conclusion, such manipulation eliminated the shift in body posture and movement of the foreleg in decapitated flies (Movie S5) indicating the efficiency of tsh-GAL80 in blocking the expression of CsChrimson. In addition, no apparent effect was observed on the backward walking of GH146^II^-, NP225-, and NP5288-GAL4 intact flies when tsh-GAL80 was expressed (Figure 4F and Movie S5). Taken together, the combined results of the above experiments suggest that GH146^II^, NP225-, and NP5288-GAL4 neurons in the central brain contribute to the backward walking we observed. These neurons function upstream and in parallel to MDNs.

**Figure 4:**
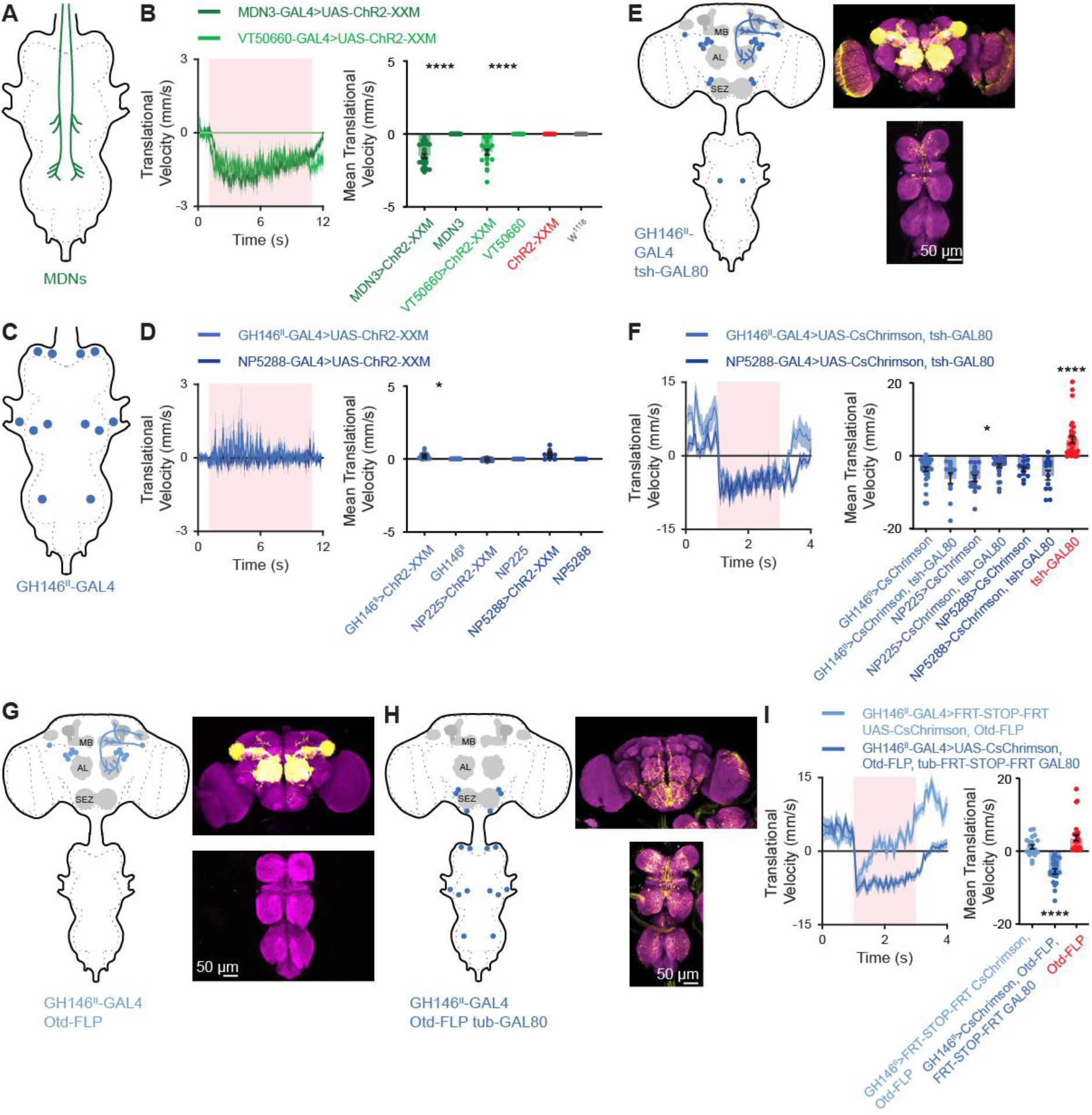
SEZ neurons underlie GH146^II^-, NP225- and NP5288-GAL4 optogenetically induced backward locomotion. **(A)** Schematic drawing of the expression pattern for the flies used in B. **(B)** Decapitated flies show backward locomotion in the case of MDN driver lines as designated. ChR2-XXM was used to activate the neurons. *Left*, translational velocity ± SEM (shading). The 10 second light pulse is labeled in red. *Right*, mean translational velocity during the light pulse obtained from flies such as presented on the left for MDN driver lines driving ChR2-XXM, parental controls, and *wt* flies. (28≥n≥12, **** p < 0.0001 for all comparisons to respective parental controls and *wt*, Kruskal – Wallis test followed by Dunn’s post-hoc test). **(C)** Schematic drawing of the expression pattern for the flies used in D. **(D)** Decapitated flies show no backward locomotion in the case of GH146^II^-, NP225- and NP5288-GAL4 driver lines. ChR2-XXM was used to activate the neurons. *Left*, translational velocity ± SEM (shading). The 10 second light pulse is labeled in red. *Right*, mean translational velocity during the light pulse obtained from flies such as presented on the left for GH146^II^-, NP225- and NP5288-GAL4 driver lines driving ChR2-XXM, and parental controls. (29≥n9, * p < 0.05, Kruskal – Wallis test followed by Dunn’s post-hoc test). **(E)** *Left,* schematic drawing of the expression pattern for the flies used in F. *Right,* expression pattern of GH146^II^-GAL4 driving UAS-CsChrimson-mVenus in the presence of tsh-GAL80 in an adult fly. Maximum intensity projection of 106 confocal sections (1 μm) through the central brain and ventral nerve cord are presented. **(F)** Blocking expression in the TAG does not affect backward locomotion in the case of GH146^II^-, NP225- and NP5288-GAL4 driver lines. CsChrimson was used to activate the neurons. *Left*, translational velocity ± SEM (shading). The 2 second light pulse is labeled in red. *Right*, mean translational velocity during the light pulse obtained from flies such as presented on the left for GH146^II^-, NP225- and NP5288-GAL4 driving CsChrimson, or CsChrimson along with tsh-GAL80 which blocks expression in the TAG. Blocking TAG expression does not abolish backward walking (33≥n≥12, * p < 0.05, **** P < 0.0001, Kruskal – Wallis test followed by Dunn’s post-hoc test). **(G)** *Left,* schematic drawing of the expression pattern for flies used in I. *Right,* expression pattern of GH146^II^-GAL4 driving UAS-FRT-STOP-FRT-CsChrimson-mVenus in the presence of Otd-nls:FLPo in an adult fly. Maximum intensity projection of 117 confocal sections (1 μm) through the central brain and ventral nerve cord are presented. **(H).** *Left,* schematic drawing of the expression pattern for flies used in I. *Right,* expression pattern of GH146^II^-GAL4 driving UAS-CsChrimson-mVenus in the presence of Otd-nls: FLPo and tub-FRT-STOP-FRT-GAL80 in an adult fly. Maximum intensity projection of 145 confocal sections (1 μm) through the central brain and ventral nerve cord are presented. **(I)** Blocking expression in the SEZ neurons affects backward locomotion in GH146^II^-GAL4. CsChrimson was used to activate the neurons. *Left*, translational velocity ± SEM (shading). The 2 second light pulse is labeled in red. *Right*, mean translational velocity during the light pulse obtained from flies as presented on the left for GH146^II^-GAL4 driving UAS-FRT-STOP-FRT-CsChrimson in the presence of Otd-nls:FLPo (G) or UAS-CsChrimson in the presence of both Otd-nls:FLPo and tub-FRT-STOP-FRT-GAL80 (H), and for Otd-nls:FLPo control. Activation of SEZ neurons led to significant backward locomotion compared to control flies. (28≥n≥21, **** p < 0.0001, Kruskal – Wallis test followed by Dunn’s post-hoc test).

### GH146^II^-_1_ NP225- and NP5288 SEZ neurons are required for backward walking

The anatomical analysis of the GH146^11^-, NP225-, and NP5288-GAL4 driver lines (Figure S3) revealed two common brain regions, the AL and the SEZ. In order to isolate in which of these brain regions the neuron or neurons which trigger the robust backward walking reside, we took advantage of the Otd-nls:FLPo transgene that expresses flippase (FLP) specifically in the brain^42^ with the exception of the gnathal ganglia in the SEZ^43^. Due to the strong similarity between GH146^II^-, NP225- and NP5288-GAL4 expression patterns (Figures S3), for this set of experiments we only used the GH146^II^-GAL4 driver line. Otd-nls:FLPo was used along with GH146^II^-GAL4 and UAS-FRT-STOP-FRT-CsChrimson to drive activity mostly in PNs (Figure 4G). Flies expressing CsChrimson in this restricted manner showed no backward walking, except for a minor transient response at the onset of the light pulse (Figure 4I). In a complementary experiment, we used Otd-nls:FLPo, GH146^II^-GAL4, UAS-CsChrimson and tub-FRT-STOP-FRT-GAL80 in order to drive expression selectively in the SEZ and the TAG, by expressing the GAL4 suppressor GAL80^44^ in the entire central brain except for the gnathal ganglia (Figure 4H, right). Expression of CsChrimson in this subset resulted in robust and sustained backward walking (Figure 4I). Recalling that neither TAG neurons nor ventromedial SEZ neurons are required for backward walking (Figure 4E, F), the combined results point to lateral SEZ neurons as underlying the sustained backward walking observed in GH146^II^-, NP225- and NP5288-GAL4 flies.

### A single SEZ neuron in each brain hemisphere drives backward locomotion

To determine which of the SEZ neurons triggers backward walking, we used the recently-developed SPARC method^45^. By expressing nSyb-PhiC31 recombinase pan-neuronally together with each of the three available SPARC2-CsChrimson::tdTomato variants (s-sparse, i-intermediate, d-dense) and GH146^II^-GAL4, we generated flies that expressed CsChrimson in different fractions of GH146^II^ labeled neurons in a stochastically distributed manner. Individual flies were then subjected to behavior experiments followed by dissection and staining to determine which neurons were labeled. We defined eight expression clusters which also included single SEZ neurons. We performed a multiple linear regression analysis (n = 85) in which the flies’ average backward walking distance was regressed against the eight defined independent variables^20^. Only a single cluster, containing a single lateral SEZ neuron in each brain hemisphere, correlated significantly with the backward locomotion observed (Figure 5A, B, S4, and Table S1). We name these two neurons, MooSEZs, for Moonwalker SEZ neurons.

**Figure 5:**
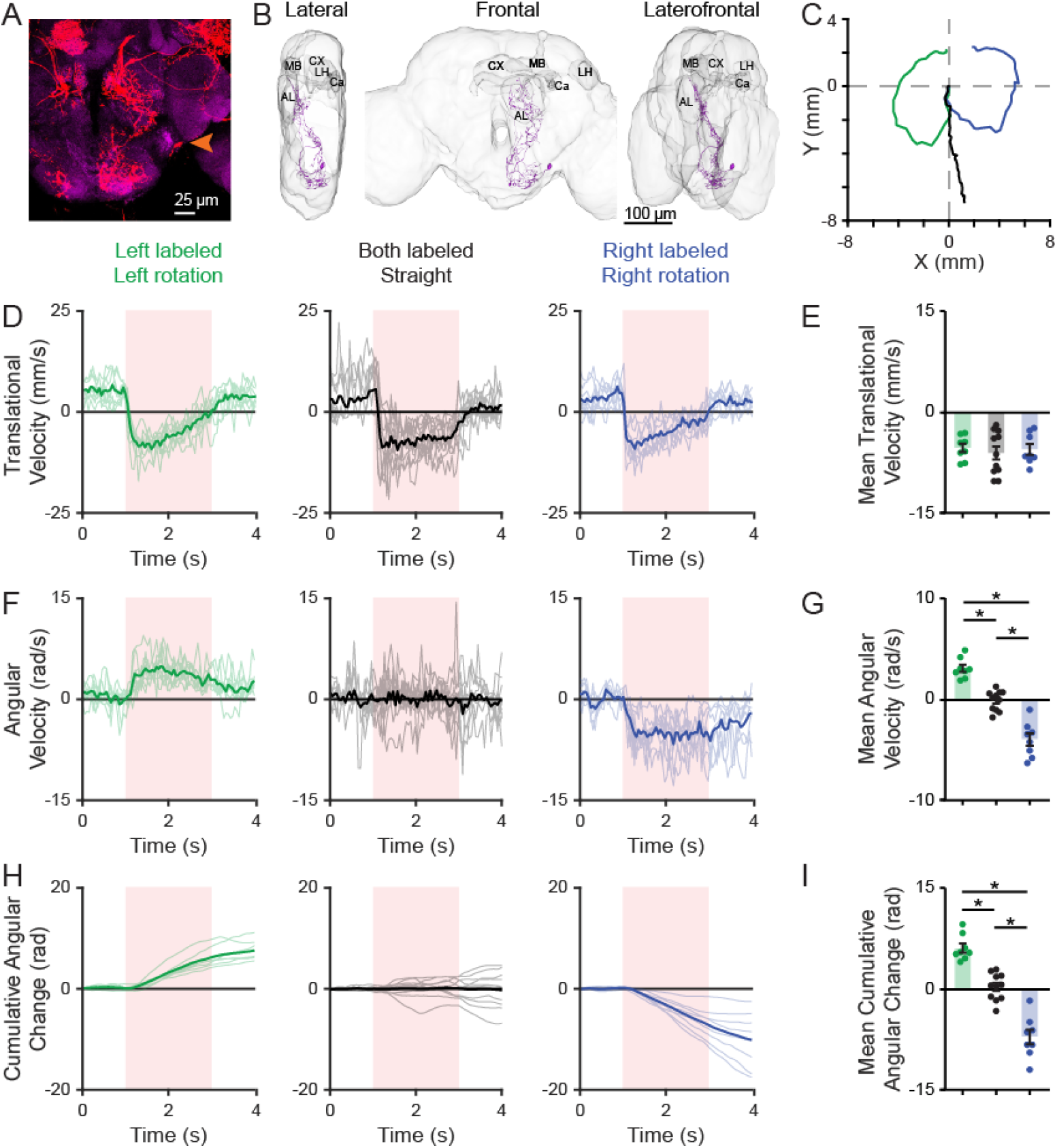
A single SEZ neuron, MooSEZ, underlies GH146^II^-, NP225- and NP5288-GAL4 optogenetically-induced backward locomotion. **(A)** Example of SPARC stochastic labeling which covered only the left MooSEZ neuron. Arrow head labels the soma. Maximum intensity projection of 70 confocal sections (1 μm) through ventral part of the central brain. **(B)** Three dimensional reconstruction of a single MooSEZ neuron. Lateral (left), frontal (middle) and laterofrontal (left) views are presented. MB, mushroom body; CX, central complex; AL, antennal lobe; Ca, calyx; LH, lateral horn. **(C)** Example of the trajectory of flies in which MooSEZs are optogenetically activated unilaterally (right – blue, left – green) or bilaterally (black). **(D, F, H)** Single fly traces (light) and mean (dark) of translational velocity (C), angular velocity (E), and cumulative angular change (G), for flies in which MooSEZs are optogenetically activated unilaterally (right – blue, left – green) or bilaterally (black). **(E, G, I)** Mean translational velocity (D), mean angular velocity (F), and mean cumulative angular change (H), for flies presented in D,F,H. Whereas all flies share similar translational velocity, the angular velocity is tightly correlated with MooSEZ side of activation. (11≥n≥8, * p < 0.05, Kruskal – Wallis test followed by Dunn’s post-hoc test).

The stochastic labelling often labeled a single MooSEZ in only one brain hemisphere (Figure 5A). This enabled examination of whether unilateral activation of MooSEZ, leads to backward rotation. Indeed, contrary to the straight backward locomotion following bilateral activation of MooSEZs, asymmetric activation of MooSEZ for 2 seconds consistently generated backward turning originating from contralateral activation (with respect to the location of MooSEZ soma in the brain) without affecting the overall backward distance passed (Figure 5 C-I). Having unequivocally identified MooSEZ as underlying GH146^II^-GAL4 mediated backward locomotion we reconstructed its structure. MooSEZ morphology implies dendritic regions mostly in the SEZ, and putative presynaptic projections in the lower lateral accessory lobe (LAL)^46^ located posterior to the AL and extending lateroventrally^47^ – a major input region of MDNs^19^. The putative presynaptic arborizations are organized in a characteristic 2-dimensional vertical sheet that slopes from anterolateral to posteriomedial (Figure 5B). Taken together, we identify in each brain hemisphere a single SEZ neuron, MooSEZ that drives contralateral activity. Combined with the results above which show clear activation of MDNs by GH146^II^-GAL4 and an MDN-independent pathway, these results suggest that MooSEZ act both upstream and in parallel to MDNs to evoke backward walking.

### An intersection approach to narrow down GH146^II^-, NP225- and NP5288-GAL4 expression pattern

The results thus far demonstrate that aversive odors can trigger backward locomotion mediated by MDNs and that MooSEZs can also trigger moonwalking when activated whether via MDNs or through a parallel pathway. However, the physiological stimulus which activates MooSEZs is not clear. GH146^II^-, NP225- and NP5288-GAL are all broad labelling driver lines, which are unsuitable for behavioral and SEZ functional imaging experiments. To try and limit the number of neurons covered by these driver lines, we used another driver line which should have a strong overlap with these three driver lines. It is well known that different transcription systems (i.e. GAL4^48^, LexA^49^, QF^50^) or genomic insertion sites, may affect the expression pattern^51,52^. We therefore examined to what extent GH146-QF recapitulates the labeled cells in GH146^II^-GAL4, which revealed notable exceptions in the SEZ (Figure 6A).

**Figure 6:**
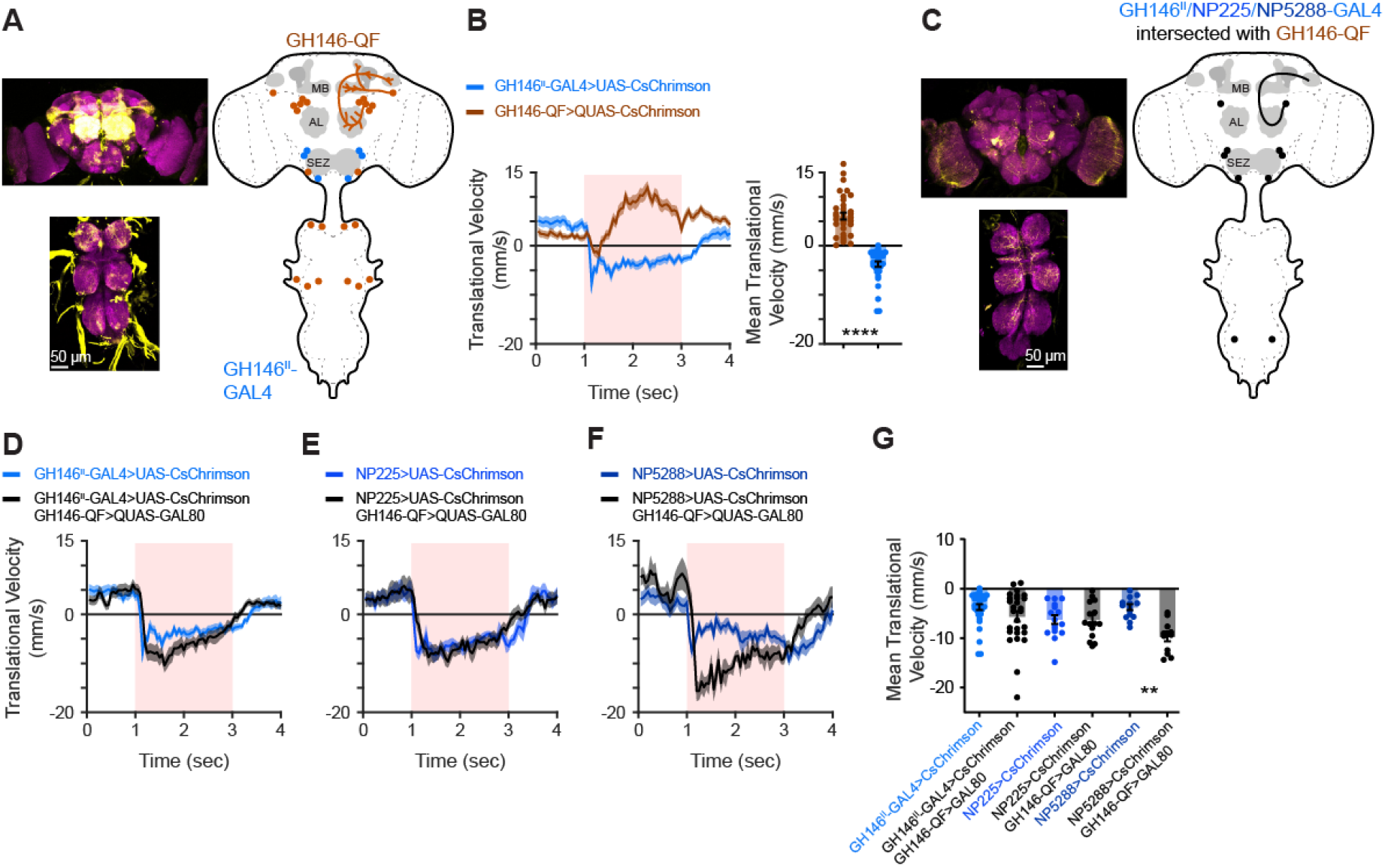
A genetic intersection approach to narrow GH146^II^-, NP225- and NP5288 expression pattern. **(A)** *Left,* expression pattern of GH146-QF driving QUAS-GFP. Maximum intensity projection of 150 confocal sections (1 μm) through the central brain and TAG are presented. *Right,* schematic drawing of the expression pattern for GH146-QF. **(B)** *Left,* translational velocity ± SEM (shading) for GH146^II^-GAL4 and GH146-QF driving CsChrimson as designated. Backward walking is observed only for GH146^II^-GAL4. Light pulse is labelled by light red. *Right,* mean translational velocity during the light pulse obtained from flies as presented on the left (n=33, **** p < 0.0001, Mann-Whitney test). **(C)** *Left,* expression pattern of an intersection between GH146^II^-GAL4 driving CsChrimson-mVenus and GH146-QF driving the GAL4 inhibitor QUAS-GAL80. Maximum intensity projection of 128 confocal sections (1 μm) through the central brain and TAG are presented. *Right*, schematic drawing of the expression pattern for GH146^II^-, NP225-, and NP5288-GAL4 when intersected with GH146-QF. **(D-F)** Translational velocity ± SEM (shading) for GH146^II^- (E), NP225- (F), and NP5288-GAL4 (G) driving UAS-CsChrimson (blue) or when intersected with GH146-QF driving QUAS-GAL80 (black). Backward walking is maintained in intersection flies and even enhanced (NP5288-GAL4). Light pulse is labelled by light red. (**G**) Mean translational velocity during the entire light pulse obtained from traces in D-F. No significant difference is observed between the backward walking velocities generated by intersection flies compared to non-intersection flies, or there is even a significant increase (NP5288-GAL4). (33≥n≥11, ** p < 0.01, Kruskal – Wallis test followed by Dunn’s post-hoc test).

The anatomical analysis suggested that GH146-QF does not overlap with GH146^II^-, NP225- and NP5288-GAL4 in the SEZ neuron relevant for backward walking. Indeed, optogenetic activation with CsChrimson of GH146-QF did not elicit backward walking (Figure 6B). We then performed an intersection between GH146^II^-, NP225- and NP5288-GAL4 with GH146-QF. GH146^II^-, NP225- and NP5288-GAL4 were used to drive CsChrimson and GH146-QF was used to drive QUAS-GAL80. Subtracting GH146-QF from GH146^II^-GAL4 strongly decreased the expression in the TAG except for a pair of ascending abdominal cells that projected towards the ventrolateral protocerebrum (Figure 6C). In addition, AL labeling was mostly abolished. In all brains examined we observed that only a single PN, which labels glomerulus VL2a, was reliably expressed (Figure S5). This PN receives input from neurons expressing the ionotropic receptor 84a (IR84a) which couples food presence to the activation of the courtship circuitry and increases male courtship^53^. In most cases we observed stochastic labeling of few other PNs, however, these were not reliably labeled across brains (Figure S5). As expected, the expression pattern of SEZ neurons was unaffected (Figure 6C). Optogenetic activation using CsChrimson of all three GAL4 drivers with subtracted GH146-QF resulted in clear and sustained backward walking (Figure 6D-G). Importantly, contrary to the stochastic nature of AL glomeruli labeling, all flies reliably performed backward walking (Figure 6G). Thus, these results further support our conclusion that MooSEZs underlie this sustained backward walking. In some cases (GH146^II^- and NP5288-GAL4), we actually observed a stronger and smoother backward walking phenotype (Figure 6D, F, G). This may arise from reduced appetitive olfactory input due to the reduced expression in AL, as activating all ORNs using ORCO-GAL4 resulted in a forward movement (Figure 2A, B), or, more likely, from the reduced foreleg activity we observed (Movie S2) which was totally abolished in these flies (Movie S6). Indeed, NP5288-GAL4 flies showed the strongest foreleg movement (Movie S7) and also showed the largest change in backward walking following the intersection (Figure 6F, G). Taken together, our intersection approach dramatically reduces the expression pattern by GH146^II^-, NP225- and NP5288-GAL4 and maintains expression in MooSEZs.

### MooSEZs Respond to Odors

We now turn to examine the physiological input that activates MooSEZs. The SEZ receives mostly gustatory and mechanosensory input^54^. Therefore, we examined these two modalities. For an aversive gustatory input we used quinine. We generated flies carrying GCaMP6f^35^ using the intersection approach and performed 2-photon *in vivo* Ca^2+^ imaging. Application of quinine did not elicit any Ca^2+^ response in MooSEZs (Figure 7A). To examine mechansensory input we used a previously described approach^20^. As before we performed 2-photon *in vivo* Ca^2+^ imaging in SEZ neurons but this time while optogenetically activating neurons that express the mechanosensory channel NOMPC ^55,56^ with CsChrimson^20^. We did not observe any Ca^2+^ response in SEZ neurons following the activation of NOMPC-expressing mechanosensory neurons (Figure 7B).

**Figure 7:**
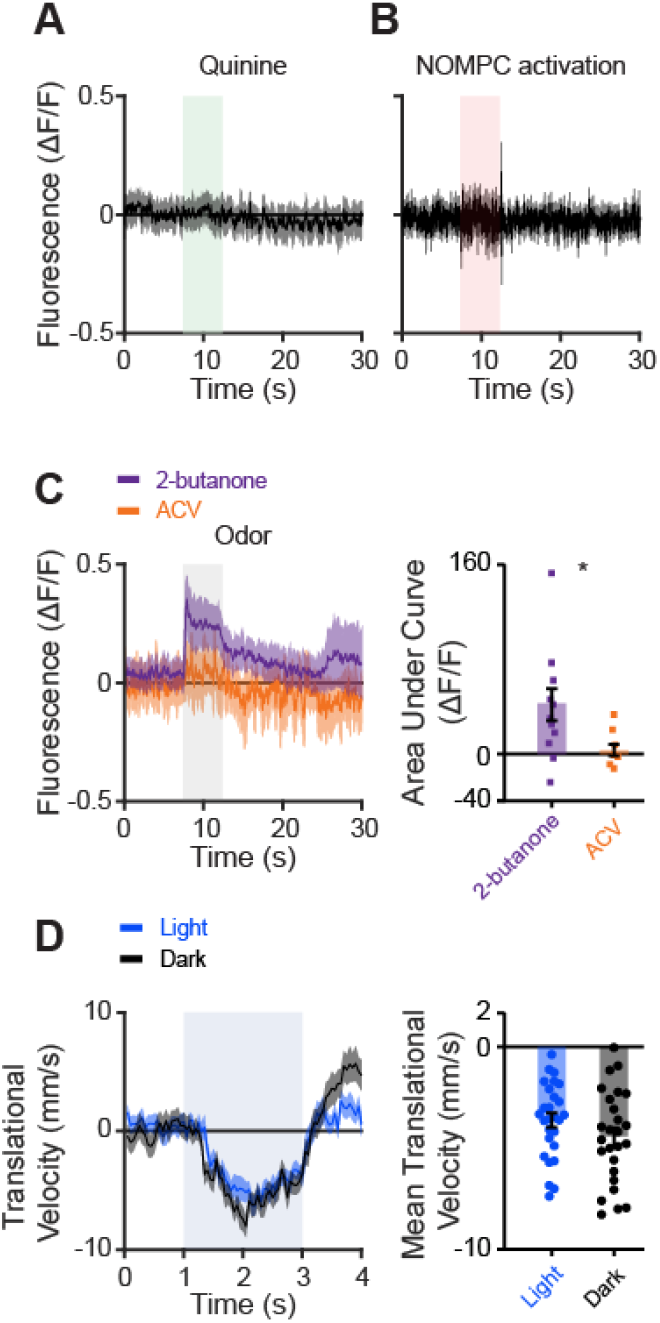
MooSEZs respond to odors. **(A)** Averaged traces ± SEM (shading) of responses following quinine application (labelled in light green). GH146^II^-GAL4 was used to drive GCaMP6f and GH146-QF was used to drive GAL80 to limit the expression pattern as presented in Figure 6. No response to quinine was observed (n=8 flies). **(B)** Averaged traces ± SEM (shading) of responses following activation of neurons expressing the mechanical sensor NOMPC (labelled in light red). NOMPC-LexA was used to drive LexAop-CsChrimson and GH146^II^-GAL4 was used to drive GCaMP6f. No response to activation of NOMPC expressing neurons was observed (n=6 flies). **(C)** *Left,* averaged traces ± SEM (shading) of responses following an odor pulse (as indicated, odor pulse is labelled in grey). GH146^II^-GAL4 was used to drive GCaMP6f and GH146-QF was used to drive GAL80 to limit the expression pattern as presented in Figure 6. *Right,* area under the curve of ΔF/F during odor response for the traces presented in the *left* panel. Similar to MDNs (Figure 1), the aversive odor 2-butanone has a stronger odor response compared to the appetitive odor ACV. (12≥n≥8, * p < 0.05, Two-sample t-test). **(D)** *Left,* translational velocity ± SEM (shading) following application of 2-butanone when MooSEZs were blocked using blue light activation of GtACR2 (35.33 mW/cm^2^) or in control experiments performed in the dark. The odor pulse is labelled in grey. GH146^II^-GAL4 was used to drive GtACR2 and GH146-QF was used to drive GAL80 to limit the expression pattern as presented in Figure 6. *Right,* mean translational velocity during the entire odor pulse obtained from traces on the left. No significant difference was observed between the backward walking velocity generated by 2-butanone when MooSEZs were active or inactive (27≥n≥26, Mann-Whitney test).

The lack of gustatory or mechanosensory signal in MooSEZs has led us to also examine olfactory input. Surprisingly, we observed a small but significant Ca^2+^ signal following application of the aversive odor 2-butanone (Figure 7C) that elicited backward walking (Figure 1E, F), but not following the more attractive odor ACV (Figure 7C) that did not elicit significant backward walking (Figure 1E, F). Since a single PN was consistently labelled in GH146^II^-GAL4 with subtracted GH146-QF flies, we repeated this experiment using the approach described above (Figure 4I) in which we used Otd-nls:FLPo, GH146^II^-GAL4, UAS-GCaMP6f and tub-FRT-STOP-FRT-GAL80 to get expression only in SEZ neurons. Similar results were obtained (Figure S6A). Thus, the cumulative results demonstrate that an aversive olfactory input arrives to MooSEZs.

The fact that odors activate MooSEZs in a specific manner and that specific optogenetic activation of MooSEZs results in backward locomotion, indicates that MooSEZs are sufficient to drive odor-driven backward locomotion. To examine if MooSEZs are also necessary for odor-driven backward locomotion we expressed in MooSEZs using the intersection approach the anion channelrhodopsins GtACR2^57^. Silencing MooSEZs had little or no effect on odor-driven backward locomotion (Figure 7D). To verify that GtACR2 efficiently blocked MooSEZs, we coexpressed it with CsChrimson under the control of GH146^II^-GAL4. Such manipulation blocked the backward walking of the flies normally elicited by the activation of CsChrimson (Figure S6B). Thus, MooSEZs are sufficient but not necessary for odor-elicited backward locomotion.

### MooSEZs drive backward crawling in larvae

Although the downstream neural architecture supporting locomotion is fundamentally different between the limbless larvae and the six-legged adult fly, MDNs preserve their function in both larvae and adult flies^58^. We therefore examined whether this is also true for MooSEZ. Anatomical analysis in larvae revealed that GH146^II^-, NP225- and NP5288-GAL4 all have expression in SEZ neurons in addition to AL and TAG (Figure S7A). In agreement with the anatomical analysis, optogenetically activating the neurons covered by these driver lines elicited backward crawling (Figure S7B and Movie S8). Surprisingly, activating MDNs using the MDN3-GAL4 driver line did not elicit backward crawling (Figure S7B). Since MDNs were demonstrated to drive backward crawling using a different driver line^58^ (R53F07-GAL4), this result suggests that MDN3-GAL4 drives expression in MDNs at a later stage of development. We then examined the expression pattern following subtraction of GH146-QF from GH146^II^-GAL4. Behavior analysis of these larvae revealed that the backward crawling was maintained. A recent study in larvae reported the existence of two SEZ neurons, AMB neurons, which mediate backward crawling via activation of MDNs^59^. In contrast, MooSEZs seem to be able to act in an MDN-independent manner. Anatomical analysis reveals great similarity between MooSEZ and AMB neurons; however, it remains to be shown if the neurons are identical or belong to different cells in the same cluster.

## Discussion

In the present study we show that aversive olfactory stimulus can trigger backward locomotion which is mediated by MDNs. We identified a single SEZ neuron in each hemisphere which triggers contralateral activity and named it Moonwalker SEZ neuron (MooSEZ). We showed that MooSEZs act both upstream and in parallel to MDNs. Although located in the SEZ, MooSEZs do not respond to the typical SEZ inputs, i.e. gustatory and mechanosensory inputs. Rather, MooSEZs respond to odors in a differential manner; they respond to an aversive odor but not to an appetitive one. Finally, we showed that MooSEZs are sufficient but not necessary for olfactory-driven backward locomotion (Figure 8).

**Figure 8:**
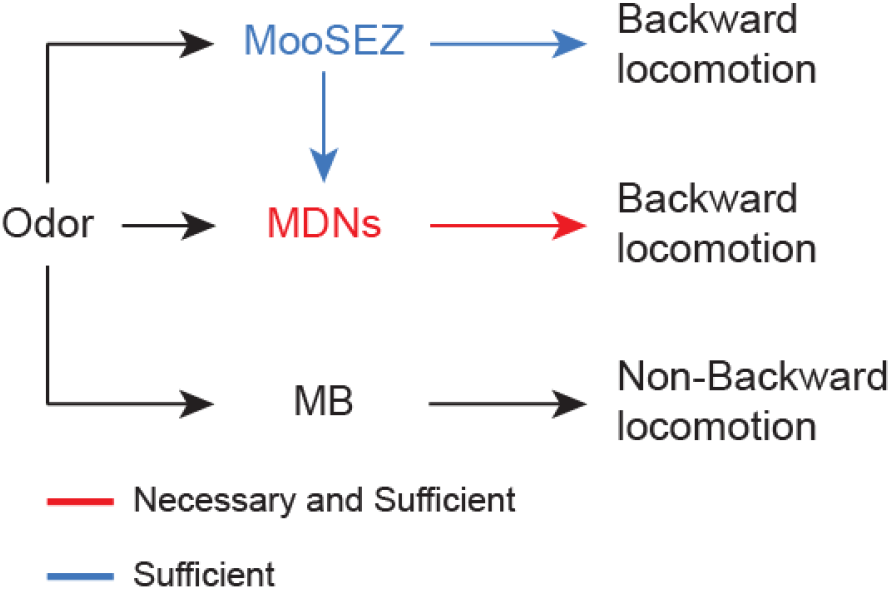
A circuit diagram of adult olfactory pathways that induce backward locomotion. Activation of the well-known MB olfactory pathway is presumably not involved in backward walking asactivation of MB neurons did not elicit backward locomotion. Odors activate MDNs and this activation is necessary for odor induced backward locomotion. Backward walking following optogenetic activation of MDNs indicates also MDNs sufficiency. Odors also activate MooSEZs. Backward walking following optogenetic activation of MooSEZs indicates MooSEZ sufficiency. However, MooSEZ are not necessary for backward locomotion as blocking them does not impair it.

The anatomy of the MooSEZ suggests dendritic regions mostly in the ventral SEZ, and putative presynaptic projections in the inferior protocerebrum. A neuron with similar behavior and morphology in larvae expresses pre- and postsynaptic markers in the respective regions (AMB^59^), supporting this notion. In adults, the protocerebral innervation is in the lower LAL, posterior to the AL, possibly reaching into the wedge, lateroventral to the AL. Since MDNs have their dendrites in the lower LAL^18,19^, a direct input from MooSEZs to MDNs is feasible in this region.

It is well established that the SEZ processes mechanosensory and gustatory sensory input^54^. In addition, it is well established that SEZ output controls movements involved in feeding behavior^60,61^. Although the putative dendritic region of MooSEZs is in the ventral SEZ, surprisingly they do not respond to bitter taste (Figure 7A) or optogenetic activation of mechanosensors (Figure 7B). Rather, MooSEZs respond to aversive olfactory input (Figure 7C). Since olfactory activity was never attributed to the SEZ it is not clear how MooSEZ receive their aversion-selective olfactory input. In larvae, anatomical studies have previously demonstrated the existence of SEZ neurons with dendritic arborizations in the AL and axonal projections in SEZ. These neurons receive specific inputs from some ORNs and from some AL local neurons^62^. In addition, AL local neurons were demonstrated to innervate both the antennal lobe and the suboesophageal ganglion (SOG)^62–64^. Multiglomerular PNs (mPNs) were demonstrated to have dendritic arborizations in SEZ, but not axonal projections^62,64^. Contrary to the larval olfactory system in the adult olfactory system local neurons do not arborize in the SOG. Only two candidate cells per hemisphere are described that could potentially convey olfactory information directly from the AL to the SEZ: lPN4 and AST3^31^. The lPN4 sends projections to the ventrolateral protocerebrum, the antennal mechanosensory and motor center (AMMC) and the medial SEZ, whereas the AST3 exclusively innervates the primary gustatory center in the medial SEZ. The lPN4 has arborizations in glomerulus VA1d and VA1v, as well as non-glomerular arborizations in the posterior AL, and a putative role in pheromone processing. The AST3 exhibits sparse multiglomerular varicose arborizations in both ALs and hence could integrate a putative olfactory warning signal arising from aversive odor blends^31^. The two second order olfactory neuropils, the LH and the MB, have either no direct connections to the SEZ^65^, or are not directly involved in the backward walking pathway described here (Figure 2A). It is therefore possible that MooSEZs receive olfactory information in a highly preprocessed way through third- or fourth-order connections, that there are additional presynaptic sites in the LAL, closer to the putative output region, or that there are other unmapped AL inputs that can convey olfactory information to the SEZ. It remains to be shown via which circuit aversive odors elicit activity in the MooSEZs.

MDNs were demonstrated to receive sensory information from the fly’s visual system through indirect synaptic connections^19^. Similarly MDNs receive mechanosensory input from the VNC^20^ but whether directly or indirectly was not addressed. Whether other sensory inputs converge onto MDNs was not resolved yet. Our results add olfaction to the multi-modality input of MDNs (Figure 1). In addition, we identify the SEZ as participating in MDN input (Figure 3A, C) and surprisingly in olfactory MDN input (Figure 7C). Our results combined with recent results in larvae^59^ suggest that SEZ neurons also act as modality integrators responding at least to olfactory and visual inputs although it still remains to be examined whether AMBs and MooSEZs are identical neurons and whether the adult MooSEZs also respond to light. In this context it seems that the mechanosensory input from the VNC bypasses the SEZ and projects directly to the region where MDN dendrites arborize^20^. The participation of SEZ as an input source to MDNs also raises the option that gustatory input, whether taste or texture, can activate MDNs.

It is to be noted that similarly to MDNs^58^, MooSEZs survive metamorphosis and persist to adulthood, to perform a functionally equivalent role of backward retreat, although the downstream neural architecture supporting locomotion is fundamentally different between the limbless larvae and the six-legged adult fly. This striking conservation through all developmental stages of this two distinct neurons, which share similarities in functionality, suggest that MDNs and MooSEZs are part of an important motor circuitry crucial for survival.

AMB neurons were described in larvae to be directly upstream to MDNs and having synaptic connections with them. In addition, MDNs seemed to be necessary for backward crawling mediated by larval AMBs. In contrast, we unraveled a parallel pathway to MDNs and recruitment of MDNs by MooSEZ could only be achieved at high stimulation intensity (Figure 3). Since our experimental approach using both dim and strong light intensities enabled us to reveal these differences, there is either a genuine difference between the architecture of the larval and adult circuits involved in MooSEZ-mediated backward locomotion, or MooSEZs and AMBs are non-identical cells in the same cluster with overlapping and somewhat complementary functionality. In this context, other experimental results are in line with the notion that there is an MDN-independent neural circuit in adult flies that contributes to backward locomotion. For instance, optogenetic activation of TLAs while blocking MDNs^20^ resulted in a residual backward walking component. In addition, backward movement was observed in association with a large membrane voltage difference between two non-MDN descending neurons^46^. Taken together, it seems that rather than a single command circuit originating in MDNs, there is at least one additional neural circuit the can mediate backward locomotion.

MDNs seem to be required for MooSEZ-driven backward locomotion primarily following strong activation of MooSEZs (Figure 3). In this context the silencing of MDNs does not result in only forward locomotion as would be expected, but rather flies fell on their back. This raises a possibility that following a strong input which drives backward locomotion, MDNs are required to maintain regular locomotion and in their absence, flies fail to walk and lose their balance.

MDNs were demonstrated to have a contralateral effect such that activation of MDNs in only one brain hemisphere led to rotation rather than straight backward locomotion^18,19^. We observe the same thing for MooSEZ unilateral activation (Figure 5). Importantly, since we used light intensities which do not result in the recruitment of MDNs (Figure 3 and S2), this reflects an MDN-independent behavior.

Our study demonstrates that an aversive olfactory cue can trigger backward locomotion via two different pathways (Figure 8). The relatively well known MDN pathway which we demonstrated to be necessary for odor evoked-backward locomotion, at least in high odor intensities, and a novel pathway which acts both upstream and in parallel to MDNs but seems to be only sufficient but not necessary to odor-driven backward locomotion (Figure 7). The existence of two parallel pathways raises the question of how one of them can be a necessary pathway. One possible option is that, contrary to the optogenetic activation of MooSEZs which generates a robust and constant activation, odor stimulation elicits a physiologically adequate response which is used for initiation of backward locomotion or supports backward locomotion, but cannot sustain backward locomotion by itself.

## Acknowledgments

We thank Dr. Robert Kittel, Dr. Christopher J Potter, Dr. Barry Dickson, Dr. Adam Claridge Chang, Dr. Andrew Lin, Dr. Gero Miesenböck, Dr. Christian Wegener, Dr. Stephen F. Goodwin, Dr. Galit Shohat-Ophir, the Bloomington Stock Center, the Vienna *Drosophila* Resource Center and the Kyoto *Drosophila* Genetic Resource Center for fly strains. This work was supported by the European Research Council (676844, MP) and the Deutsche Forschungsgemeinschaft (HU 2474/1-1, WH).

## Author contributions

SI: Initiated the project, conceptualization, methodology, investigation, formal analysis, software, writing–review & editing, visualization. ER: Investigation, formal analysis, visualization, writing–review & editing. DW: Investigation, visualization. WH: Investigation, visualization, writing–review & editing. MP: Initiated the project, conceptualization, methodology, investigation, formal analysis, writing–original draft, writing– review & editing, visualization, supervision, funding acquisition.

## Methods

### LEAD CONTACT AND MATERIALS AVAILABILITY

Further information and requests for resources and reagents should be directed to and will be fulfilled by the Lead Contact, Moshe Parnas.

Flies generated for this paper, data and code used to generate the figures will be available upon request.

### EXPERIMENTAL MODEL AND SUBJECT DETAIL

#### Fly Strains

Fly strains (see below) were raised on cornmeal agar under a 12 h light/12 h dark cycle at 25 °C. The following transgenes were used: GH146^II^-GAL4 (Bloomington #30026^66^), GH146^X^-GAL4^67^, GH146-QF (Bloomington #30037, #30038^50^), MDN3-GAL4^18^, w^1118^ (Bloomington # 5905), Or82a-GAL4 (Bloomington #23125), Ir31a-GAL4 (Bloomington #41726), Ir40a-GAL4 (Bloomington #41727), Ir25a-GAL4 (Bloomington #41728), Ir76b-GAL4 (Bloomington #41730), Ir8a-GAL4 (Bloomington #41731), Ir64a-GAL4 (Bloomington #41732), Ir92a-GAL4 (Bloomington #41733), Ir84a-GAL4 (Bloomington #41734), Ir76a-GAL4 (Bloomington #41735), GMR22C06-GAL4 (Bloomington #48974), GMR24G07-GAL4 (Bloomington #49095), GMR68D02-GAL4 (Bloomington #39471), GMR26B04-GAL4 (Bloomington #49158), GMR48E05-GAL4 (Bloomington #50370), GMR24A10-GAL4 (Bloomington #49059), GMR95C02-GAL4 (Bloomington #48431) NP6115-GAL4, NP3062-GAL4, NP5021-GAL4, NP5103-GAL4, NP80-GAL4, NP2001-GAL4, NP2297-GAL4, NP3529-GAL4, NP5288-GAL4, NP225-GAL4, NP1579-GAL4, Orco-GAL4 (Bloomington #26818^68^), VT50660-GAL4^18^, VT040053-GAL4’ VT019428-GAL4, VT026020-GAL4, MZ612-GAL4^69^, MB247-GAL4 (Bloomington #50742^70^), c305a-GAL4 (Bloomington #30829^71^), OK107-GAL4 (Bloomington #854^72^), LN1-GAL4 (KYOTO Stock Center #103945^73^), LN2-GAL4 (KYOTO Stock Center #104198^73^), Mz699-GAL4^74,75^, LC16-1-GAL4 (Bloomington #68331^19^), TLA-GAL4^20^ (Bloomington #74205, Bloomington #72912), MDN-LexA, UAS-CsChrimson-mCherry^19^, nSyb-PhiC31 (Bloomington #84151^45^), SPARC2-D-CsChrimson::tdTomato (Bloomington # 84143^45^), SPARC2-I-CsChrimson::tdTomato (Bloomington # 84144^45^), SPARC2-S-CsChrimson::tdTomato (Bloomington # 84145^45^), UAS-CsChrimson (Bloomington #55135, #55136^37^), UAS-ChR2-XXM^76^, UAS-GtACR2^57^, UAS-TNT (Bloomington #28838^36^), UAS-TNT inactive (Bloomington #28839^36^), UAS-FRT-STOP-FRT-CsChrimson-mVenus, tub-FRT-STOP-FRT-GAL80 (Bloomington, #39213), NOMPC-LexA (Bloomington #52241^55,56^). LexAop-CsChrimson-mVenus (Bloomington, #55138), UAS>dsFRT>CsChrimson-mVenus, UAS-GCaMP6f (Bloomington #42747, #42748, and #52869^35^), UAS-mCD8-GFP (Bloomington #32185), LexAop-mCD8-GFP (Bloomington #32203), LexAop-GCaMP6m-p10 (Bloomington #44275, #44276), Otd-nls:FLPo, QUAS-GAL80, tsh-GAL80, LexAop-TNT, QUAS-CsChrimson#32c

For activation of CsChrimson or GtACR2, flies were collected 2-5 days post eclosion and grown for another 3-5 days on 1mM *all-trans* retinal (R2500; Sigma-Aldrich) supplemented food in complete darkness before experimental testing was performed. When starved flies were used, they were placed in a fresh vial containing water-soaked filter paper 24 hours prior to the experiment.

#### Behavioral Analysis

##### Adult flies

Experiments were conducted in two behavioral chambers: an open arena^77^ of 60mm diameter and linear chambers^18^, 40*1.5*1.5 mm Length\width\height mm long (8 linear grooves in a plate). Both chambers were composed of polyoxymethylene and covered with a transparent acrylic plastic. For the open arena, odor (2-butanone or ACV) was applied through a 5mm diameter hole in the plastic lid located at the center of the open arena, while for the linear chambers odor was delivered through a 5mm hole that was further connected by a narrow air stream path to one end of the linear chambers. Flies were allowed to walk freely for 1-3 minutes before start of an experimental trial.

For optogenetic experiments, flies were illuminated with either 617nm LED (M617L3; THORLABS) for CsChrimson activation, or with 470nm LED (M470L3; THORLABS) for ChR2-XXM or GtACR2 activation. For olfactory experiments, a 2 sec odor pulse (2-butanone or ACV) was delivered. Odors at 5*10^-2^ final dilution were delivered by switching mass-flow controlled carrier and stimulus streams at a final flow of 0.8 l/min (CMOSens Performance Line, Sensirion) via software controlled solenoid valves (The Lee Company) through a 1/16 inch ultra-chemical-resistant Versilon PVC tubing (Saint-Gobain, NJ, USA). 2-butanone was diluted in mineral oil (Sigma-Aldrich, Rehovot) and ACV in DDW. Odors were prepared on a daily basis. Throughout performed experiments, the open arena and linear chambers were illuminated from the bottom by a high intensity 810nm IR LED (SFH 4786S; OSRAM), while Flies’ behavioral responses were recorded from the top by a camera (PL-D795MU-T; PIXELINK), equipped with 16mm focal length lens (NMV-16M11; NAVITAR) coupled to a 800nm long-pass filter (LP800; MIDOPT), at 20 frames per second and 832×832 pixel resolution.

In open arena experiments, groups of 3-15 female and male flies were loaded, and presented with either a 2 or 10 second light stimulus, or a 2 second odor pulse, except for SPARC^45^ stochastic activation experiments, in which female flies were tested individually and illuminated with a 2 second light pulse for 3 consecutive trials. In linear chamber experiments, 8 flies were loaded in 8 separate grooves in a plate and were exposed to a 2 second odor stimulus.

##### Larvae

Male and female third instar larvae that were kept in a complete darkness throughout development were collected and transferred to a 1% agarose arena with a diameter of 85mm. Larvae were allowed to crawl freely for 1-3 minutes before the start of an experimental trial, in which a 10sec blue (470 nm, activating ChR2-XXM) or red (617 nm, activating CsChrimson) was applied. When CsChrimson was used larvae were grown on 1mM all-*trans* retinal (R2500; Sigma-Aldrich) supplemented food in complete darkness. Groups of 3-8 larvae were placed in the arena for a single trial that consisted a 10 second light pulse. In the case of intersection between GH146^II^ and GH146-QF, each larva was tested individually.

##### Video analysis

Acquired videos were tracked with Ctrax^78^ which allows to determine the locations and orientations of flies along a recorded video. Following initial tracking with Ctrax, the “FixErrors Matlab GUI” was used to compensate for tracking errors. Only flies that were accurately tracked were considered in the analysis. The “BehavioralMicroarray Matlab Toolbox” was used to compute per-frame parameters of the flies’ (and larvae’s) locomotor activity. Custom MATLAB scripts were used to perform final analysis based on the parameters outputted by “The “BehavioralMicroarray Matlab Toolbox”. Translational velocity was defined as the projection of the fly’s velocity vector on its forwardbackward orientation axis. Angular velocity was defined as the change in the fly’s orientation angle in relation to a global orientation coordinate system. Backward walking distance was computed by integrating the area under negative translational velocity against time and cumulative angular change was computed by integrating the area under the angular velocity against time. Fraction of time of total backward locomotion was computed by dividing the number of time frames in which flies were measured with negative translational velocity during the light pulse by the number of time frames in which light was delivered. Percentage of backward walking flies was calculated by defining a minimal backward walking distance of 3mm as a threshold for walking activity that had a pronounced backward component during the light stimulation. Accordingly, backward walking percentage was calculated as the ratio between the number of flies that walked backwards 3mm and above to the total number of tested flies multiplied by 100. For Regression analysis of SPARC^45^ stochastic flies, each cluster was set to 1 if at least one cell within the cluster was targeted, and to 0 in case in which none of the cells within the cluster were labeled.

##### Functional Imaging

Functional imaging was performed on 5-10 days post-eclosion females using a two-photon laser-scanning microscopy (DF-Scope installed on an Olympus BX51WI microscope) as previously described^34,79,80^. The brain was superfused with carbonated solution (95% O_2_, 5% CO_2_) containing 103 mM NaCl, 3 mM KCl, 5 mM trehalose, 10 mM glucose, 26 mM NaHCO_3_, 1 mM NaH_2_PO_4_, 3 mM CaCl_2_, 4 mM MgCl_2_, 5 mM N-Tris (TES), pH 7.3. Odors at 5*10^-2^ final dilution were delivered by switching mass-flow controlled carrier and stimulus streams at a final flow of 0.8 l/min (Sensirion) via software controlled solenoid valves (The Lee Company). Air-streamed odor was delivered through a 1/16 inch ultra-chemical-resistant Versilon PVC tubing (Saint-Gobain, NJ, USA) that was placed 5 mm from the fly’s antenna. Fluorescence was excited by a Ti-Sapphire laser (Mai Tai HP DS, 100 fs pulses) centered at 910 nm, attenuated by a Pockels cell (Conoptics) and coupled to a galvo-resonant scanner. Excitation light was focussed by a 20X, 1.0 NA objective (Olympus XLUMPLFLN20XW), and emitted photons were detected by GaAsP photomultiplier tubes (Hamamatsu Photonics, H10770PA-40SEL), whose currents were amplified (Hamamatsu HC-130-INV) and transferred to the imaging computer (MScan 2.3.01). All imaging experiments were acquired at 30 Hz. When necessary, movies were motion-corrected using the TurboReg ^81^ ImageJ plugin. ΔF/F was calculated as previously described^34,79,80^.

##### Structural Imaging

Brain dissections, fixation, and immunostaining were performed as described^34,80^. To visualize native GFP, tdTomato and mVenus fluorescence in both larval and adult brains, we followed our previous adult brain protocol^82^. In short, dissected brains were fixed in 4% (w/v) paraformaldehyde in PBS (1.86 mM NaH_2_PO_4_, 8.41 mM Na_2_HPO_4_, 175 mM NaCl) and fixed for additional 20 min at room temperature. Samples were washed for 3×20 min in PBS containing 0.3% (v/v) Triton-X-100 (PBT) and blocked with 5% normal goat serum for 30 min. Samples were then incubated for 2 days with primary antibodies. After being washed three additional times with PBT, samples were incubated for 2 days with secondary antibodies in PBT followed by embedding in Vectashield. Primary antibodies used: rabbit anti-GFP (Thermo Fisher Scientific A-11122, 1:1000), mouse anti-GFP (Abcam ab1218, 1:250), rabbit anti-RFP (Abcam ab62341, 1:250), mouse anti-Bruchpilot (DSHB, 1:50). Secondary antibodies used: Alexa Flour 488 goat anti-mouse (Abcam ab150113, 1:500), Alexa Flour 488 goat anti-rabbit (Thermo Fisher Scientific A-11034, 1:500), Alexa Flour 568 goat antirabbit (Abcam ab175471, 1:500), Alexa Flour 647 goat anti-mouse (Abcam ab150115, 1:500). For Larval brains no antibodies were used. Larval brains were mounted on poly-L-lysin-covered coverslips, adult brains were mounted between tape spacers. Images were collected on a Leica TCS SP5, SP8, Zeiss LSM 800, or Nikon A1 confocal microscope and processed in ImageJ. Neuron 3D reconstructions were done as described previously^39^.

### QUANTIFICATION AND STATISTICAL ANALYSIS

#### Statistics and data analysis

Statistical testing and parameter extraction were done using GraphPad Prism and custom MATLAB code (The MathWorks, Inc.). All statistical tests details can be found in the figure captions. Significance was defined as a p-value smaller than 0.05 and all statistical tests were two-sided, except for a (left) one-sided t-test performed in Figure 2B.

For presentation, bar plots with dots were generated using the UnivarScatter MATLAB ToolBox (https://www.mathworks.com/matlabcentral/fileexchange/54243-univarscatter) and the shadedErrorBar function (https://github.com/raacampbell/shadedErrorBar) for shaded errors on imaging traces.

### DATA AND CODE AVAILABILITY

The data and code used to generate figures 1–7 and supplementary figures S1-S7 are available from the corresponding author on request.

