## Supplemental Information for "Two Parallel Pathways Mediate Olfactory-Driven Backward Locomotion"

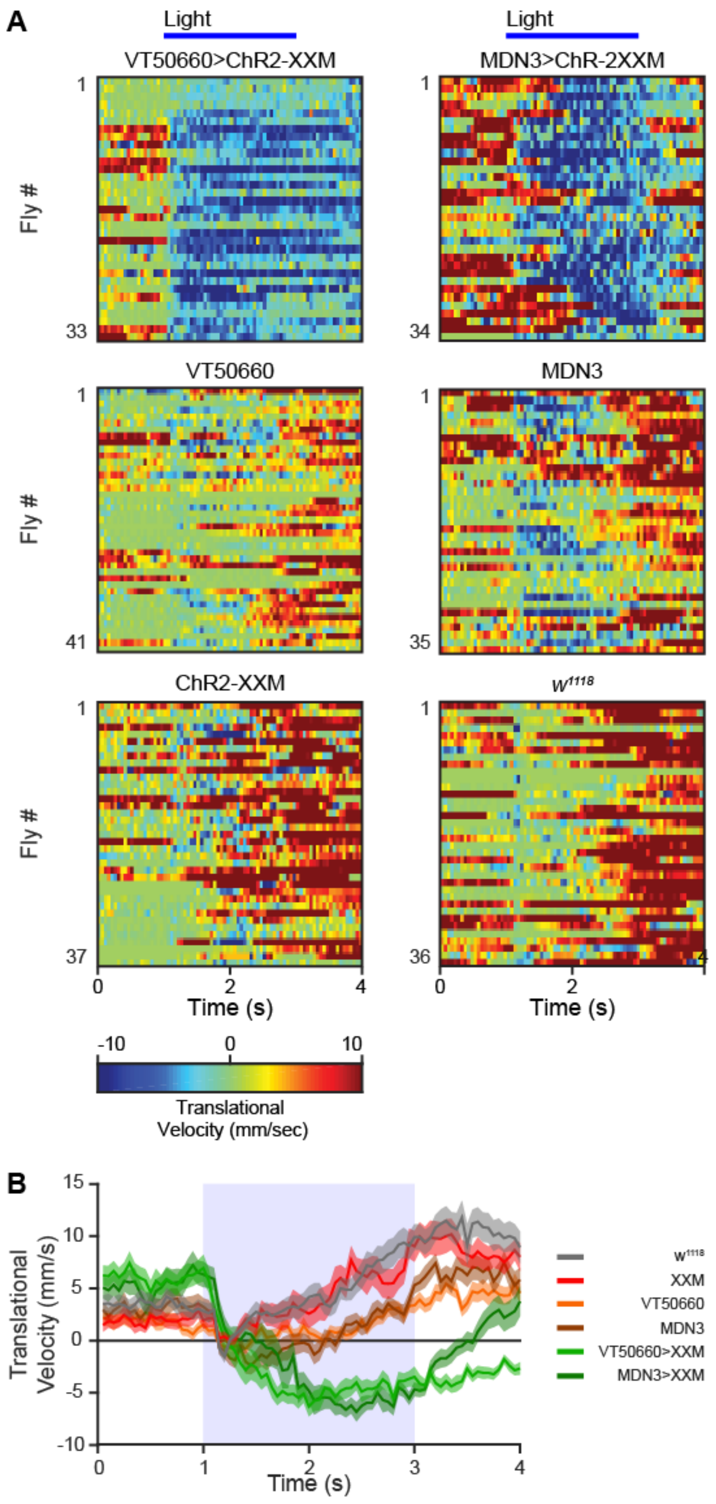


**Figure S1: Sustained backward walking following MDN activation.**
**A.** Translational velocity matrix of single flies. Genotypes as designated. A light pulse (horizontal blue line) was given between 1 and 3 seconds.
**B.** Mean translational velocity ± SEM (shading) over time for the genotypes presented in A.

Figure S2

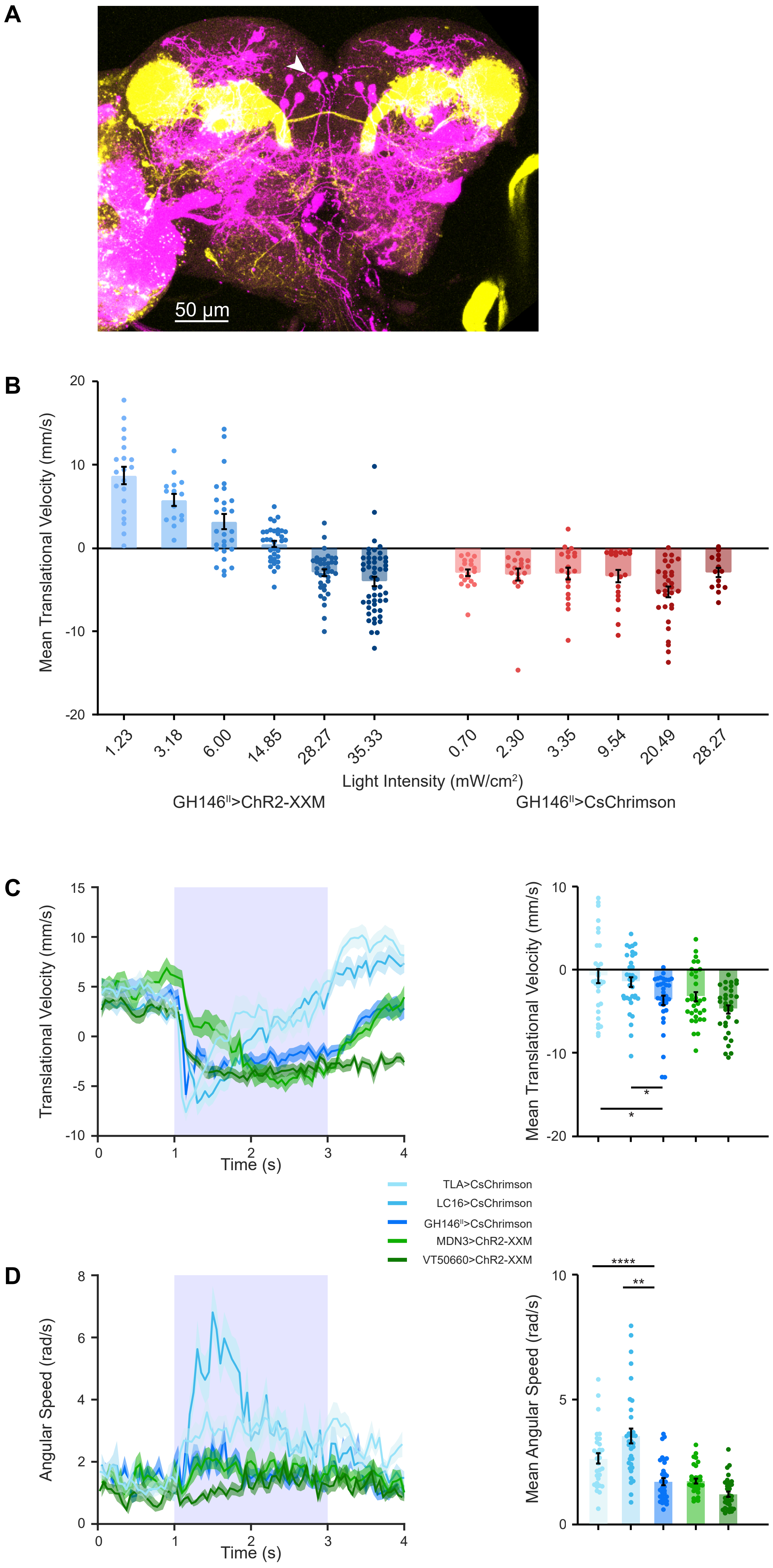

A
50 µm
B
Mean Translational Velocity (mm/s)
Light Intensity (mW/cm²)
GH146ᴵᴵ>ChR2-XXM
GH146ᴵᴵ>CsChrimson
1.23
3.18
6.00
14.85
28.27
35.33
0.70
2.30
3.35
9.54
20.49
28.27
C
Translational Velocity (mm/s)
Time (s)
Mean Translational Velocity (mm/s)
TLA>CsChrimson
LC16>CsChrimson
GH146ᴵᴵ>CsChrimson
MDN3>ChR2-XXM
VT50660>ChR2-XXM
D
Angular Speed (rad/s)
Time (s)
Mean Angular Speed (rad/s)

**Figure S2: Characterization of GH146$^{II}$-GAL4 mediated backward locomotion**

**A.** GH146$^{II}$-GAL4 was used to drive CsChrimson-mCherry. MDNs were labelled using VT44845-LexA driving GFP. White arrowhead marks MDN cell bodies. No overlap is observed between GH146$^{II}$ labeled neurons and MDNs. Maximum intensity projection of 85 confocal sections (1 µm) through the central brain are presented.
**B.** Effects of light intensity on optogenetic induced backward locomotion. GH146$^{II}$-GAL4 was used to drive either CsChrimson or ChR2-XXM. The mean translational velocity during the entire light pulse obtained for different light intensities is presented.
**C, D.** *Left,* Translational velocity (C) and angular speed (D) ± SEM (shading) elicited by activation of visual projection neurons (LC16-1-GAL4), TwoLumps Ascending neurons (TLA-GAL4), GH146$^{II}$-GAL4, or MDNs (VT50660-GAL4 or MDN3-GAL4) driving either UAS-ChR2-XXM or UAS-CsChrimson as designated. Sustained backward walking is observed throughout the two second light pulse (labelled by light blue) only for MDNs and GH146$^{II}$ driver lines. *Right*, mean translational velocity (C) and mean angular speed (D) during the entire light pulse obtained from traces on the left. TLA and LC16 had significantly lower translational velocity and increased angular speed. ($34 \geq n \geq 31$, * $p < 0.05$, ** $p < 0.01$, **** $p < 0.0001$, Kruskal - Wallis test followed by Dunn's post-hoc test).

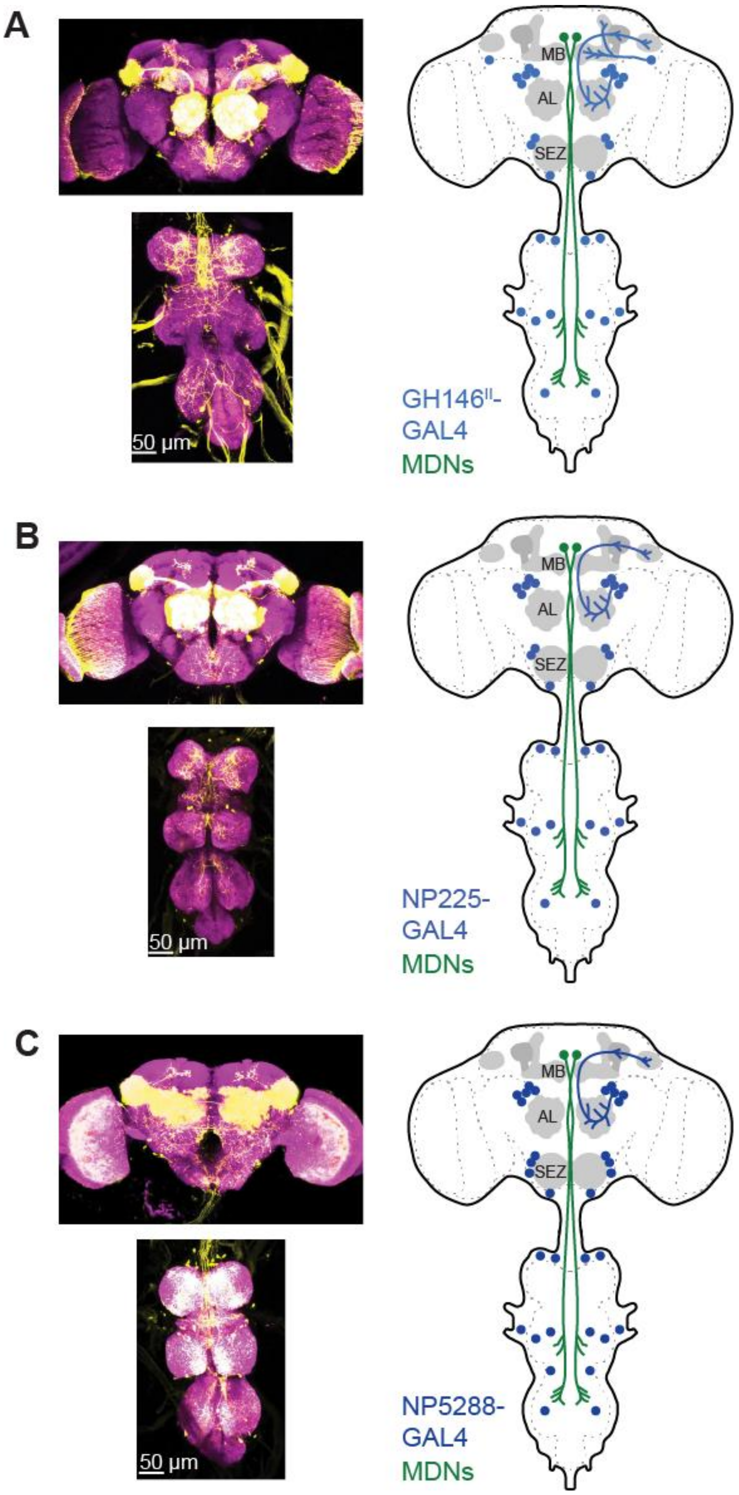


**Figure S3: GH146$^{II}$-, NP225- and NP5288-GAL4 anatomy.**

**A-C.** *Left*, Expression pattern of GH146$^{II}$-GAL4 (A), NP225-GAL4 (B), and NP5288-GAL4 (C). UAS-CsChrimson-mVenus was used to label the cells. Maximum intensity projection of 150 confocal sections (1 µm) through the central brain and TAG are presented. *Right*, Schematic drawing of the expression pattern of the driver lines on the left (blue) and of MDNs (green).

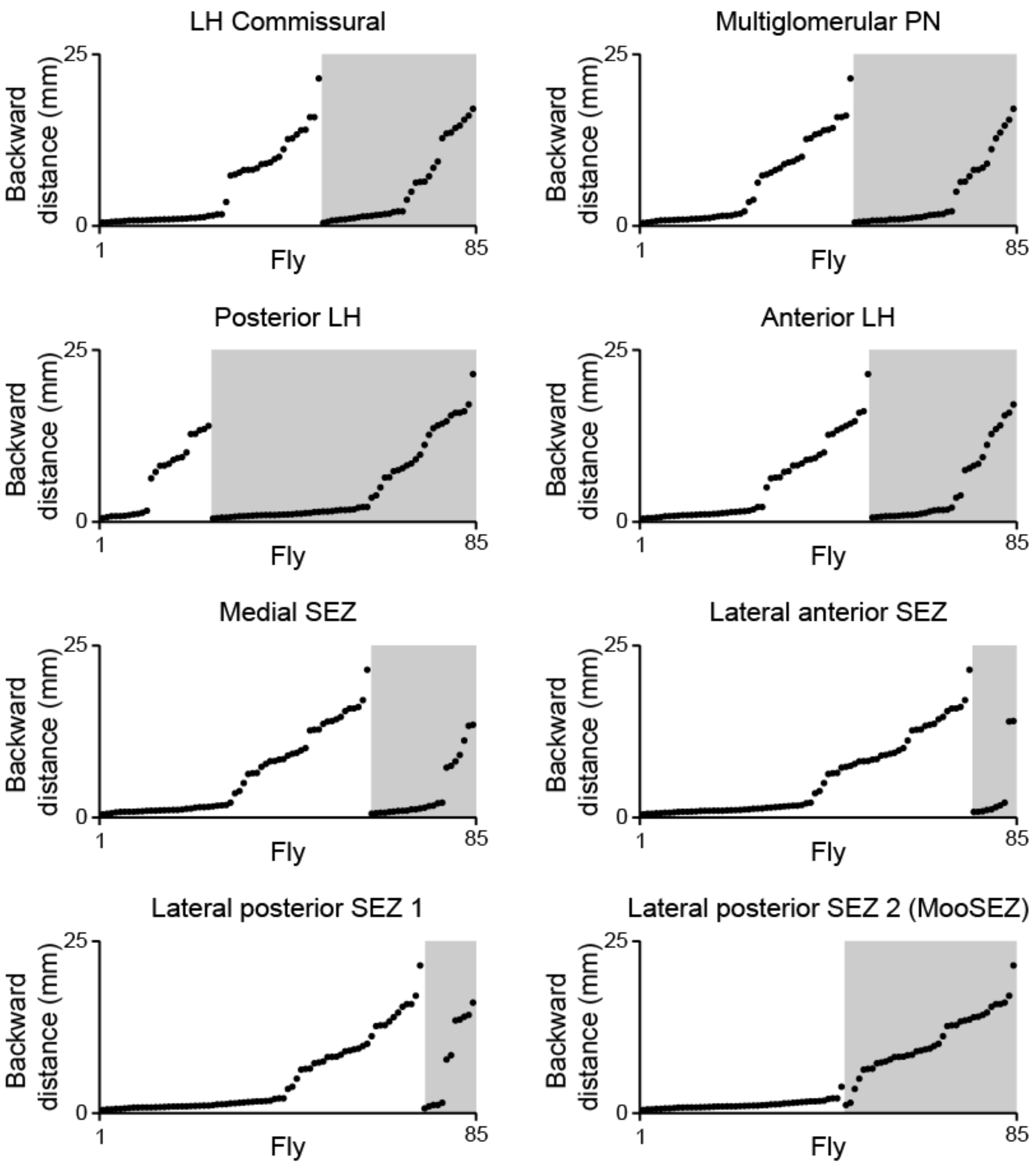


**Figure S4: A single SEZ neuron, MooSEZ, underlies GH146$^{II}$-GAL4 optogenetically-induced backward locomotion.**

Analysis of the correlation between the expression in the eight different clusters (as designated) and the mean backward locomotion distance averaged over 3 trials covered by the flies. The grey shading indicates expression in the cluster of interest. Only for MooSEZ (lateral posterior SEZ 2) a clear lack of backward walking is observed when there is no expression (white background) and a clear backward locomotion when there is expression (grey background). Multiple linear regression analysis in which each cell cluster was set as a predictor of the mean backward locomotion distance revealed a significant correlation only between the Lateral anterior SEZ 2 neuron (MooSEZ) and backward walking. (Table S1, n=85, R-squared=0.6311, Adjusted R squared=0.5922, **** $p < 0.0001$).

### Figure S5

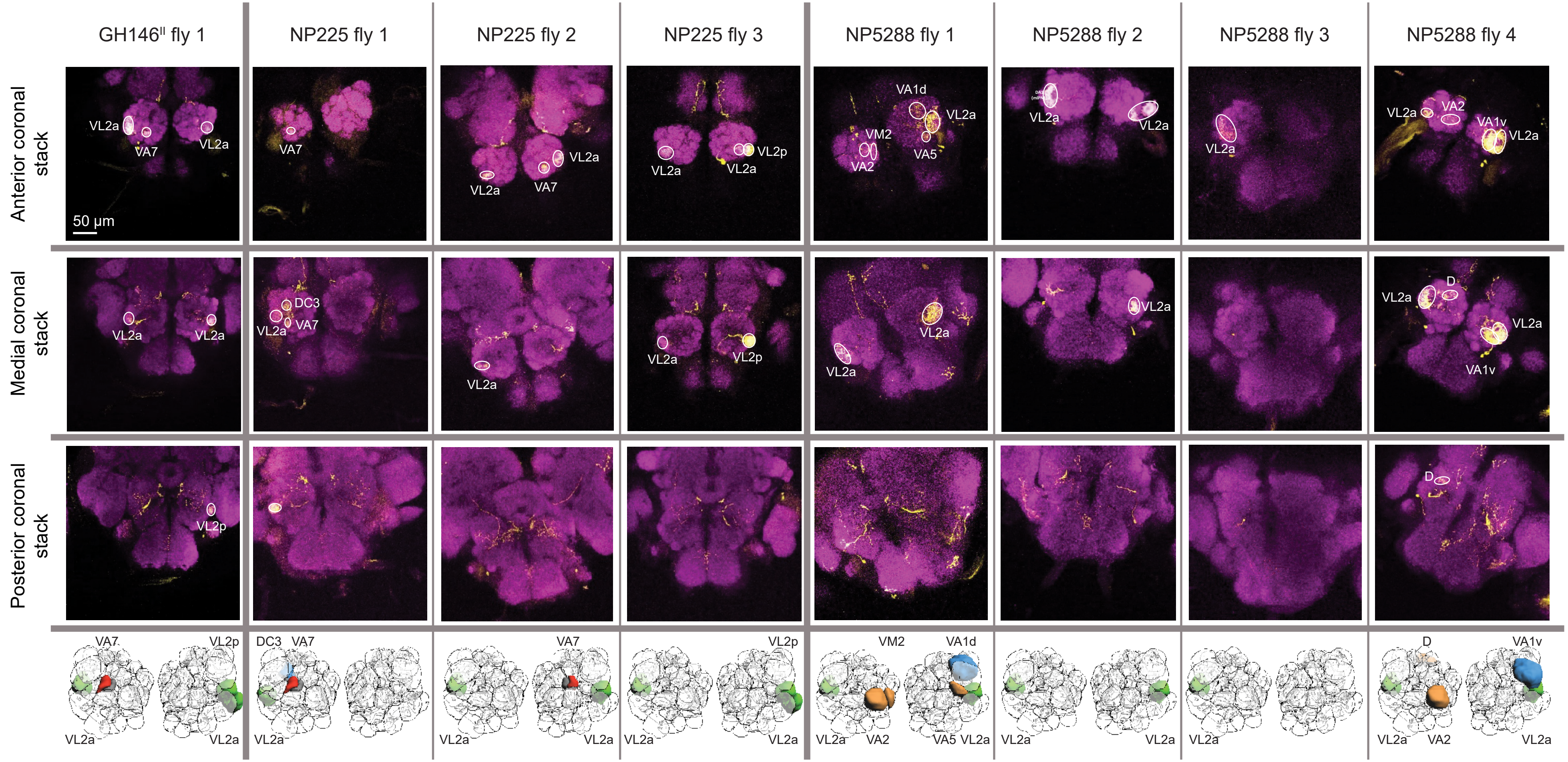

GH146[II] fly 1
NP225 fly 1
NP225 fly 2
NP225 fly 3
NP5288 fly 1
NP5288 fly 2
NP5288 fly 3
NP5288 fly 4
Anterior coronal stack
Medial coronal stack
Posterior coronal stack
50 μm
VL2a
VA7
VL2p
DC3
VA1d
VM2
VA2
VA5
VA1v
D

**Figure S5: Glomeruli labeled by intersection of GH146$^{II}$-, NP225-, NP5288-GAL4 with GH146-QF.** *Top*, AL expression patterns when GH146$^{II}$-, NP225-, or NP5288-GAL4 driving CsChrimson-mVenus were intersected with GH146-QF driving the GAL4 inhibitor QUAS-GAL80 in individual flies. Maximum intensity projections of ~20 confocal sections (1 µm) through anterior, medial and posterior coronal stacks of the antennal lobe are presented. Only a small and relatively consistent subset of AL glomeruli is labeled across different flies. *Bottom*, schematic 3D illustrations (Grabe et al., 2015) of the spatial locations of the labeled AL glomeruli.

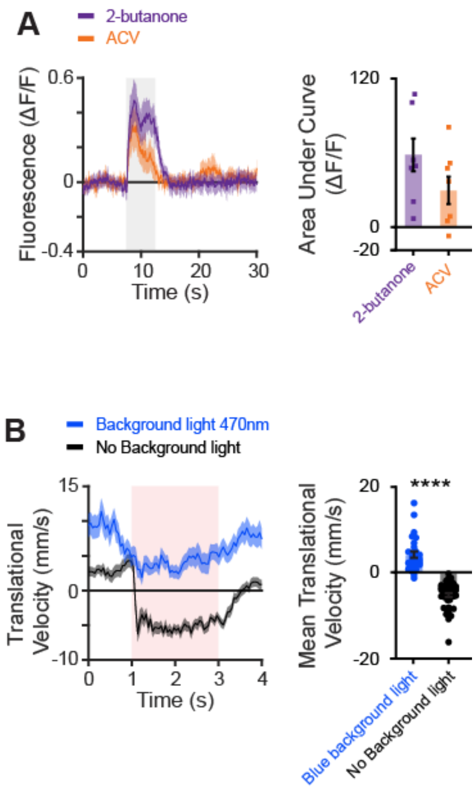


**Figure S6: GtACR2 efficiently blocks MooSEZs**

**A.** *Left,* averaged traces ± SEM (shading) of responses following an odor pulse (as indicated, odor pulse is labelled in grey). Otd-nls:FLPo, GH146[II]-GAL4, UAS-GCaMP6f and tub-FRT-STOP-FRT-GAL80 were used to drive expression of GcaMP6f in TAG and SEZ neurons. *Right,* area under the curve of ΔF/F during odor response for the traces presented in the *left* panel.

B. *Left,* translational velocity ± SEM (shading) following optogenetic activation of MooSEZ with CsChrimson (red, 3.35 mW/cm$^2$) or when the inhibitory GtACR2 channel was co-activated (blue, 35.33 mW/cm$^2$). When GtACR2 was activated the blue light (470 nm) was on through the experiment. The red light (617 nm) activating the CsChrimson was provided between the first and third seconds (red shading). GtACR2 efficiently blocks MooSEZ activation by CsChrimson. *Right*, mean translational velocity during the red light pulse obtained from traces on the left. A significant difference is observed between the backward walking velocity generated with and without the optogenetic activation of GtACR2. (45≥n≥29, **** $p < 0.0001$, Mann-Whitney test).

Figure S7

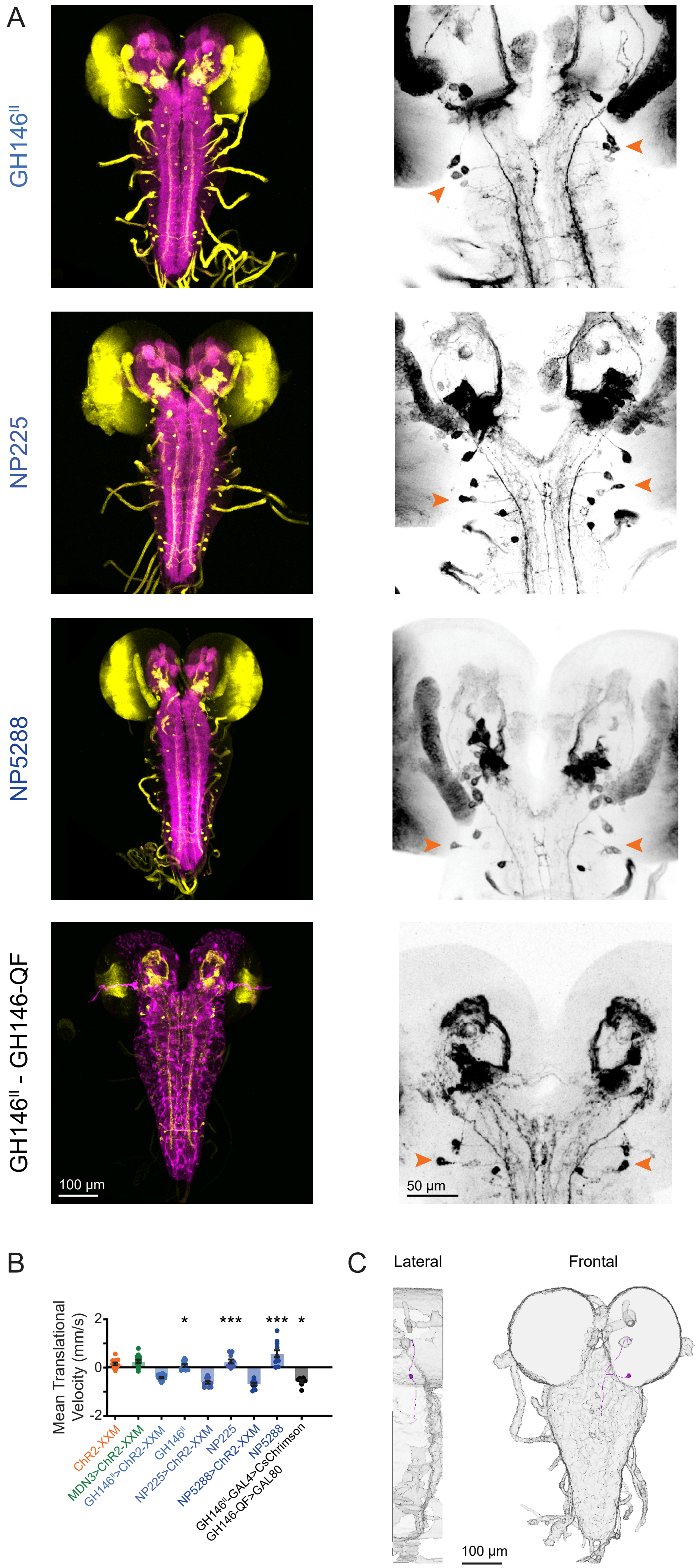
A
GH146^II
NP225
NP5288
GH146^II - GH146-QF
100 µm
50 µm
B
Mean Translational Velocity (mm/s)
2
0
-2
*
***
***
*
ChR2-XXM
MDN3>ChR2-XXM
GH146^II>ChR2-XXM
GH146^II
NP225>ChR2-XXM
NP225
NP5288>ChR2-XXM
NP5288
GH146^II-GAL4>CsChrimson
GH146-QF>GAL80
C
Lateral
Frontal
100 µm

**Figure S7: MooSEZs induce backward crawling in larvae and resemble AMBs.**
**A.** *Left,* Expression pattern in larvae of GH146$^{II}$-, NP225- NP5288-GAL4 and GH146$^{II}$-GAL4 with subtracted GH146-QF. UAS-GFP or UAS-CsChrimson-mVenus were used to label the cells. Maximum intensity projections of 60 confocal sections (1.5 µm) through the central brain and ventral nerve cord are presented. *Right,* Close-up of larval SEZ neurons labeled by GH146$^{II}$-, NP225- and NP5288-GAL4 and GH146$^{II}$-GAL4 intersected with GH146-QF. MooSEZs are designated with orange arrowheads.
**B.** Mean translational velocity of larval crawling during a 10s light pulse for the genotypes presented. ($13 \geq n \geq 8$, Pairwise comparisons with respective GAL4 parental control: * $p < 0.05$, *** $p < 0.001$, ****, Kruskal - Wallis test followed by Dunn's post-hoc).
**C.** Three dimensional reconstruction of a single larval MooSEZ.

| Variable | Parameter | Estimate | Standard error | \|t\| | p |
|---|---|---|---|---|---|
| Intercept | β0 | 2.113 | 0.9984 | 2.116 | 0.0376 |
| LH Commisural | β1 | -0.02719 | 0.8117 | 0.03349 | 0.9734 |
| Multiglomerular PN | β2 | -0.539 | 0.8249 | 0.6534 | 0.5155 |
| Posterior LH | β3 | -0.2457 | 0.8732 | 0.2814 | 0.7792 |
| Anterior LH | β4 | -0.3551 | 0.8141 | 0.4362 | 0.6639 |
| Medial SEZ | β5 | -1.323 | 0.8872 | 1.492 | 0.14 |
| Lateral anterior SEZ | β6 | -0.9135 | 1.241 | 0.736 | 0.464 |
| Lateral posterior SEZ 1 | β7 | -0.6537 | 1.183 | 0.5527 | 0.5821 |
| Lateral posterior SEZ 2 (MooSEZ) | β8 | 8.378 | 0.8227 | 10.25 | <0.0001 |

$$y \sim \beta_0 + \beta_1 X_1 + \beta_2 X_2 + \beta_3 X_3 + \beta_4 X_4 + \beta_5 X_5 + \beta_6 X_6 + \beta_7 X_7 + \beta_8 X_8$$

**Table S1: Multiple linear regression analysis for stochastic activation via the SPARC genetic method.**
**Related to figure S4**

Eight explanatory variables representing eight stochastically labeled neuronal clusters in $GH146^{II}$-GAL4 were used to predict average backward walking covered by single flies (n=85). Y denotes the dependent variable, $\beta_0$ denotes the y-intercept and $\beta_n$ denotes the slope of the $X_n$ independent variable. R squared: 0.6311, adjusted R squared: 0.5922, F(8, 76)=16.25, p<0.0001

**Movie S1: Differential odor-induced backward locomotion.**

Example of $W^{1118}$ responses to ACV (left) and 2-butanone (right).

**Movie S2: Backward locomotion following optogenetic activation of GH146$^{II}$-GAL4.**

Example of backward locomotion following a 2 sec blue light pulse (470 nm) in flies carrying GH146$^{II}$-GAL4 and ChR2-XXM.

**Movie S3: GH146-GAL4-induced, MDN independent, backward locomotion.**

*Left,* example of the efficiency of TNT in silencing MDNs. The VT50660-GAL4 driver was used to drive UAS-CsChrimson, and the VT44845-LexA (MDN-LexA) was used to drive LexAop-TNT. *Right,* example of backward locomotion generated by activation of GH146$^{II}$-GAL4 with UAS-CsChrimson while VT44845-LexA (MDN-LexA) driving LexAop-TNT was used to silence MDNs.

**Movie S4: GH146$^{II}$-GAL4 TAG neurons cannot induce backward locomotion.**

*Left,* example of backward locomotion in MDN3-GAL4 driving UAS-CsChrimson in decapitated flies. *Right,* example of lack of backward locomotion in GH146$^{II}$-GAL4 driving UAS-CsChrimson in decapitated flies. Note the movement of the forelegs.

**Movie S5: Blocking expression in TAG neurons does not impair optogenetic induced backward locomotion.**

*Left,* example of the efficiency of tsh-GAL80 to block TAG expression. No foreleg movement is observed following optogentic activation. *Right,* example of backward locomotion in NP5288-GAL4 driving UAS-CsChrimson when expression in TAG was blocked using tsh-GAL80.

**Movie S6: Backward locomotion following optogenetic activation of the neurons covered by GH146$^{II}$-GAL4 with subtracted GH146-QF.**

Example of backward locomotion following a 2 sec red light pulse (617 nm) in flies carrying GH146$^{II}$-GAL4, UAS-CsChrimson, GH146-QF and QUAS-GAL80.

**Movie S7: Blocking expression in TAG neurons enhances backward locomotion in NP5288-GAL4 flies**

*Left,* example of backward locomotion in NP5288-GAL4 driving UAS-CsChrimson. Note the strong foreleg movements. *Right,* example of backward locomotion in NP5288-GAL4 with subtracted GH146-QF driving UAS-CsChrimson. Note the absence of foreleg movement and increase in backward locomotion.

**Movie S8: Backward crawling following optogenetic activation of the neurons covered by GH146$^{II}$-GAL4 with subtracted GH146-QF.**

Example of backward crawling following a red light pulse (617 nm) in larvae carrying GH146$^{II}$-GAL4, UAS-CsChrimson, GH146-QF and QUAS-GAL80.
